# DNA 6mA marks transcriptionally active chromatin in malaria parasites

**DOI:** 10.64898/2026.06.12.732001

**Authors:** Dhanasekaran Seshan, William Lauer, Gautam Sarkar, Mohan Govindasamy, Connor Sean Murray, Eric Lieberman Greer, Melissa Laird Smith, Shruthi Sridhar Vembar

## Abstract

DNA N6-methyladenine (6mA) has emerged as a significant epigenetic modification across a broad range of eukaryotes, from unicellular protists to metazoa. However, its role in unicellular eukaryotic parasites with highly AT-rich genomes, such as malaria-causing *Plasmodium falciparum*, remains unclear. Using mass spectrometry, South-western blotting, and Single Molecule Real-Time sequencing (Pacific Biosciences) across four stages of *P. falciparum* intra-erythrocytic development (IED), we confirmed that 0.02–0.04% of genomic adenines are modified to 6mA, with over 60% of the sites being stably maintained during the IED cycle. Notably, 6mA is enriched at transcription start sites, with genes bearing 6mA marks within their 5’ and 3’ untranslated regions exhibiting significantly elevated steady-state transcript levels. Consistent with this, 6mA loci show a strong positive correlation with activating histone post-translational modifications, while showing no significant association with repressive histone marks. Furthermore, in contrast to unicellular ciliates such as *Oxytricha* and *Tetrahymena –* organisms that share ancestry with *Plasmodium –* 6mA-marked genomic regions do not occlude nucleosomes. Lastly, we identified a putative 6mA methyltransferase belonging to the METTL4 family in *P. falciparum,* PfN6AMT encoded by the *PF3D7_1303100* gene, and demonstrate that recombinant PfN6AMT exhibits robust methyltransferase activity *in vitro*, with mutation of its active site residues abolishing catalytic activity. Collectively, our findings demonstrate that 6mA is a low-abundance, yet reproducible, feature of the *P. falciparum* epigenome that is associated with transcriptionally active chromatin, and that the molecular mechanisms governing DNA adenine methylation may have undergone substantial evolutionary divergence, even among closely related eukaryotic lineages.

## INTRODUCTION

To respond to developmental and environmental cues, an organism has to rapidly vary its gene expression profile, and that too in a reversible fashion. One way to accomplish this is to fine-tune DNA accessibility and chromatin structure through epigenetic changes including (but not limited to) DNA base alterations, which is observed in both pro and eu-karyotes; histone post-translational modifications (PTMs), which is a characteristic of eukaryotes; non-coding RNAs; and nuclear architecture. In particular, small chemical tags such as methyl and acetyl groups, when added to DNA nitrogenous bases or histone tails, can control the intermolecular connections between DNA strands and chromatin machinery, thus controlling whether gene transcription is activated or silenced.

The first chemical tag to be described was methylation of cytosine at its 5^th^ carbon in DNA, 5-methylcytosine (5mC) [1]. Subsequently, 5mC was found to be abundant in the genomes of more recently evolved eukaryotes including mice and humans, and primarily repress gene expression when found within gene promoters [2], [3]. In addition to 5mC, its oxidised version, 5-hydroxymethylcytosine (5hmC), has also been identified and has been shown to activate gene expression [4], [5]. In prokaryotes, the first modified DNA base to be identified was N6-methyladenine (6mA) [6], with 5mC being detected in bacterial DNA only in the 1960s, but at significantly lower levels [7]. While both of these bases are part of DNA restriction-modification systems that protect the bacterial genome from the action of endonucleases, later studies in *Caulobacter* found that promoter adenine methylation can alter transcriptional output and cell cycle progression suggesting an epigenetic role for 6mA [8]. This was further confirmed in other bacteria including *Escherichia coli*, *Vibrio cholerae*, *Salmonella enterica*, and *Helicobacter pylori* [9], [10], [11], [12].

6mA has historically not been as well studied as 5mC presumably due to its lower abundance in more recently evolved eukaryotes [13], [14]. Although 6mA was proposed to be functional in the genome of the unicellular, free-living ciliates *Tetrahymena pyriformis* and *T. thermophila* in the early 1970s and 1980s [15], [16], it was only in the 2010s that advancements in chromatography and mass spectrometry techniques, the availability of specific anti-6mA antibodies [17], [18], and progress in nascent DNA sequencing techniques such as single-molecule real-time (SMRT) sequencing [19], enabled the robust detection of 6mA in the nuclear genomes of diverse eukaryotes including *Chlamydomonas reinhardtii* [20], *Oxytricha trifallax* [21]*, Caenorhabditis elegans* [22], *Drosophila melanogaster* [23], *Oryza sativa* [24], *Mus musculus* [13], [25] and even humans [26], [27]. In fact, the ciliate *Oxytricha trifallax* is reported to have a 6mA abundance of 0.78-1.04% in its genome [21] while early diverging fungi show up to 2.8% of their adenines to be methylated [28], which is comparable to the abundance of 5mC found in humans [29] consequently, 6mA determines nucleosome positioning in *Oxytricha* [21]. Taken together with observations linking 6mA to gene activation [20], [27], [30], transgenerational inheritance [22] and dynamic positioning during development [24], [25], [27], DNA adenine methylation has now been established as a functional epigenetic mark in eukaryotes. This has been further complemented with the identification of DNA 6mA methyltransferases that belong to the MT-A70 family including MTA1 and MTA9 in *Oxytricha* [21], the METTL4 protein DAMT-1 in *C. elegans* [22], and N6AMT1 in humans [27], [31]. Despite these efforts, the presence and epigenetic significance of 6mA in mammals remains controversial with human N6AMT1 (hN6AMT1) also possessing protein methyltransferase activity [32], [33].

The apicomplexan parasite *Plasmodium falciparum*, responsible for the most fatal form of human malaria, undergoes multiple stage transitions as it develops within the *Anopheles* mosquito and human host. These stage transitions are determined by changes in gene expression, which are regulated at the epigenetic, transcriptional, translational, and post-translational levels. Epigenetic regulation, including histone PTMs, non-coding RNAs and nuclear architecture, is particularly important during asexual intra-erythrocytic development (IED) in humans and has been linked to *P. falciparum* virulence, malaria pathogenesis and sexual commitment [34], [35], [36]. While several studies have attempted to study the role of DNA base modifications in parasite epigenetic regulation [37], [38], [39], [40], thus far, 5mC has only been detectable at very low levels and a 5hmC-like modification has been linked to gene activation, although its exact chemical nature is remains unknown [40]. Given that *P. falciparum* has one of the most AT-rich genomes sequenced to date (approximately 80.6% A+T content) and is closely related to ciliates like *T. thermophila* and *Oxytricha*, researchers have tried to detect methylated adenine bases in the parasite genome during IED, with a 2016 study reporting 0.15% of adenines as being modified to 6mA by mass spectrometry [41]. The authors also used Dpn1-Assisted 6mA-seq (DA-6mA-seq) to determine the extent of G(6mA)TC modifications in the *P. falciparum* genome and found this to be less than 10%. Overall, while DA-6mA-seq showed promise as a method to assess genomes with low 6mA abundance, no further studies have reported a regulatory role for 6mA in *P. falciparum*.

To address this, we used mass spectrometry and SMRT sequencing, which detects 6mA presence with single nucleotide resolution, and confirmed 6mA to be present anywhere from 0.02% to 0.05% in the *P. falciparum* genome at four different timepoints of asexual development. Subsequent genome-wide distribution analysis revealed a stable 6mA profile across the four timepoints, with less than 20% variability. Furthermore, 6mA was found to be enriched within gene bodies, primarily within 5’ untranslated regions (UTRs), exons and 3’ UTRs, and correlated with activating histone PTMs. As a consequence, genes with 6mA in the 5’ or 3’ UTRs showed higher steady state transcript levels. Interestingly, unlike in *Oxytricha*, where 6mA occludes nucleosomes, in *P. falciparum*, 6mA is enriched in nucleosome-containing genomic regions. Lastly, we identified a candidate DNA 6mA methyltransferase PfN6AMT encoded by *PF3D7_1303100* in the parasite genome and demonstrated its 6mA methyltransferase activity *in vitro*. Taken together, our results suggest that the DNA base modification 6mA may constitute another layer of epigenetic regulation during *P. falciparum* IED.

## RESULTS

### 6mA DNA base modifications are present at low levels in the *P. falciparum* genome

To establish the presence of 6mA in the *P. falciparum* genome, we first used commercial anti-6mA antibodies in South-western dot blotting assays. A concentration gradient of genomic DNA ranging from 25 to 400 ng prepared from cultures of the *P. falciparum* 3D7 strain synchronised at the ring, trophozoite or schizont stages of IED was used for this analysis, with genomic DNA isolated from human WBCs and total RNA isolated from *P. falciparum* 3D7 mixed stages serving as positive and negative controls, respectively. As shown in **Fig. 1A**, genomic DNA of all three stages exhibited a 6mA signal that was comparable to human WBCs, with 6mA levels relatively stable across *P. falciparum* IED (**Supplementary Fig. 1**). To validate these results, high molecular weight (HMW) *P. falciparum* genomic DNA extracted from ring (8-12 hours post invasion (hpi)), trophozoite (28-32 hpi) or schizont (40-44 hpi) stages of the *P. falciparum* 3D7 strain was subjected to liquid chromatography-tandem mass spectrometric analysis (LC-MS/MS; [42]), which identified 6mA abundance to be 0.039% (±0.003%), 0.045% (±0.001%) and 0.041% (±0.002%), respectively, indicating that, of the 9,200,000 adenines present in the ∼23 Mb *P. falciparum* genome, 3,600–4,200 adenines are methylated at any given intraerythrocytic stage (**Fig. 1B**).

**FIGURE 1.**
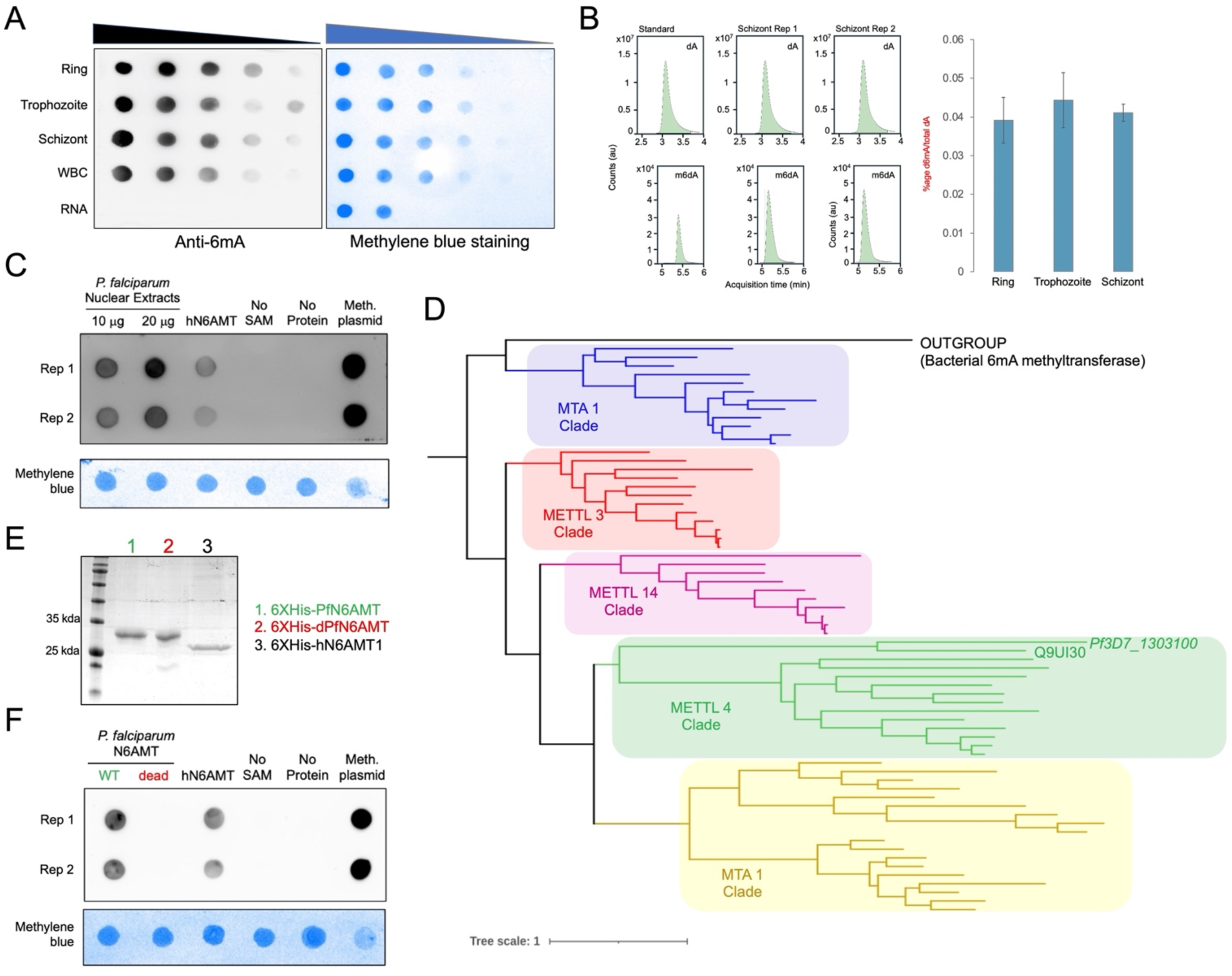
6mA was successfully detected in the genome of *P. falciparum* asexual stage parasites. ***(A)*** South-western blotting analysis of genomic DNA prepared from Ring (R), Trophozoite (T) or Schizont (S) stage parasites of the *P. falciparum* 3D7 strain using mouse monoclonal antibodies against 6mA (Synaptic Systems). Human genomic DNA prepared from WBCs and parasite RNA from a mixed stage culture served as +ve and-ve controls, respectively. Denaturation was achieved by heating the genomic DNA at 95°C for 5 min. HPI = Hours post invasion. Right panel shows the methylene blue staining of genomic DNA used*. **(B)*** Mass spectrometric detection of 6mA in *P. falciparum* 3D7 genomic DNA. ***Left panel:*** LC-MS/MS spectra for adenine (top) and 6mA (bottom). ***Right panel:*** Quantification of%6mA relative to total A content. Data represent the average ± SEM of three independent biological replicates. **(C)** 6mA methyltransferase activity assays using 10 and 20 µg of *P. falciparum* 3D7 nuclear extracts. 1 µg of 6XHis-hN6AMT1 as positive control. **(D)** Phylogenetic tree of nucleobase methyltransferase homologs from eukaryotes constructed to identify the clade that PfN6AMT (encoded by *PF3D7_1303100)* belongs to. The tree was constructed for 10,000 iterations as described by Beh *et al.*, 2019. **(E)** Coomassie-stained SDS-PAGE gel analysis of different hexahistidine (6XHis)-tagged recombinant proteins purified from bacteria using Ni^2+^-NTA affinity chromatography. **(F)** 6mA methyltransferase activity assays using 1 µg of the indicated proteins. For D and F, reactions without the methyl donor, SAM, or without protein, served as negative controls. Methylated plasmid was used as a loading control for blotting.

### 6mA methyltransferase activity can be detected in *P. falciparum* nuclear extracts

Previous studies have shown that 6mA methyltransferase activity is detectable in nuclear extracts prepared from various eukaryotic organisms [21], [43]. To demonstrate a similar property for *P. falciparum*, we extracted nuclear components from mixed stage parasites and performed *in vitro* methyltransferase activity assays using S-adenosyl methionine (SAM) as the methyl donor; recombinant hN6AMT1 [27] with an N-terminal hexahistidine tag, *i.e.,* 6xHis-hN6AMT1, served as a control. In line with our expectations, parasite nuclear extracts exhibited significant methyltransferase activity toward a synthetic oligonucleotide substrate (**Fig. 1C**), hinting at the presence of an active 6mA methyltransferase in the *P. falciparum* nucleus.

### Discovery of a putative 6mA methyltransferase PfN6AMT *(PF3D7_1303100)* in *P. falciparum*

To identify the putative *P. falciparum* 6mA methyltransferase, we retrieved the amino acid sequences of known eukaryotic 6mA methyltransferases (**Supplementary Table S1**) and performed BLASTp against the *P. falciparum* proteome. This analysis identified one possible hit when hN6AMT1 was used as query: a putative methyltransferase encoded by *PF3D7_1303100* (https://plasmodb.org). This protein, which we refer to as PfN6AMT, shares 32% sequence identity and 50.4% sequence similarity with hN6AMT1, and contains the conserved active site motif NPPY at positions 132-135 (**Supplementary Fig. S2A**). Furthermore, the AlphaFold-predicted structure of PfN6AMT aligns with the solved crystal structure of hN6AMT1 with an RMSD value of 1.6 Å (**Supplementary Fig. S2B**), with phylogenetic analysis placing both proteins in the METTL4 clade of 6mA methyltransferases (**Fig. 1D**). Lastly, to confirm that PfN6AMT possesses methyltransferase activity, recombinant hexahistidine-tagged PfN6AMT (*i.e.,* 6xHis-PfN6AMT) and a catalytically dead version 6xHis-dPfN6AMT, in which the NPPY motif was mutated to AAAA, were purified from a bacterial heterologous system (**Fig. 1E** and **Supplementary Fig. S3**) and tested *in vitro* in methyltransferase activity assays; 6xHis-hN6AMT1 served as a positive control. While recombinant 6xHis-PfN6AMT efficiently increased 6mA levels in the synthetic oligonucleotide substrate, similar to 6xHis-hN6AMT1, the catalytically dead version, 6xHis-dPfN6AMT, appeared to have lost this activity (**Fig. 1F**). Taken together, our results suggest that PfN6AMT is a putative DNA 6mA methyltransferase, which may regulate 6mA levels in the *P. falciparum* genome during IED.

### Genome-wide mapping of 6mA during *P. falciparum* IED

Subsequently, to study the genome-wide distribution pattern of 6mA during the *P. falciparum* IED, HMW genomic DNA was isolated from tightly synchronized *P. falciparum* parasite cultures of the *P. falciparum* 3D7 strain at four different IED timepoints: 10, 20, 30 and 40 hpi (three biological replicates each), and subjected to SMRT sequencing, a technique that identifies 6mA with single-nucleotide resolution in a strand-specific manner [44]. We first quantified 6mA abundance relative to the total sequenced adenine residues by considering 6mA calls with a minimum coverage of 25X, and found that the values ranged from 0.024 to 0.032% (**Fig. 2A**), and were comparable to the LC-MS/MS-derived % 6mA values, albeit 25% lower (**Fig. 1B**). Next, based on SMRT sequencing-derived genomic coordinates of 6mA, we correlated the biological replicates of a given timepoint to each other and found that they were positively correlated with a Pearson correlation coefficient value greater than 0.8; with the only outlier being replicate 3 for 20 hpi (**Fig. 2B** and **Supplementary Fig. S4**). Accordingly, for the remainder of the study, 6mA signals detected in at least two of the three biological replicates were retained as true 6mA sites (**Supplementary Table S2**).

**FIGURE 2.**
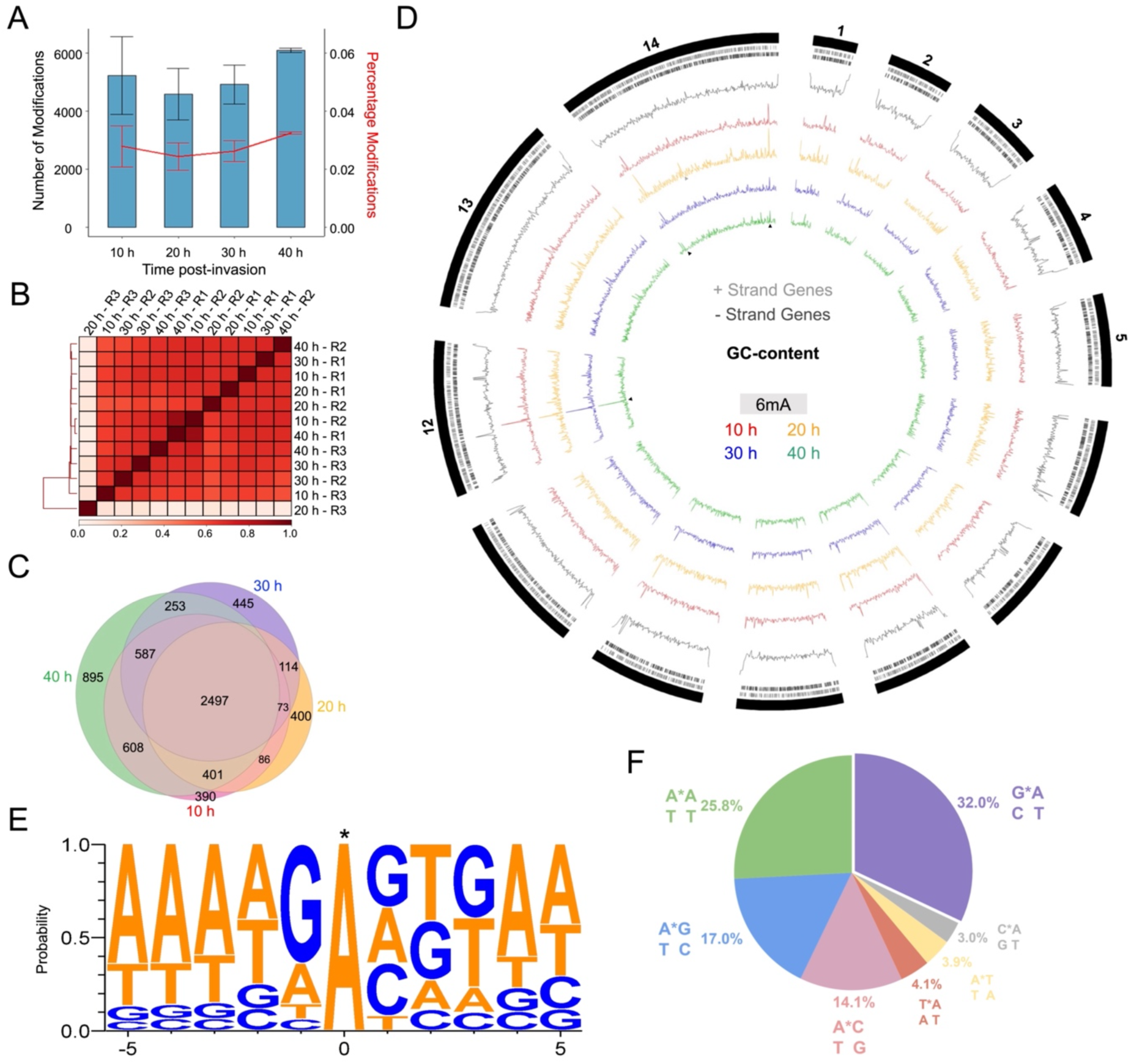
Genome-wide distribution analysis of 6mA in *P. falciparum* using SMRT sequencing. ***(A)*** SMRT sequencing-based quantification of 6mA levels in the *P. falciparum* 3D7 genome at four timepoints of asexual development. The left Y-axis indicates the average 6mA count at each timepoint while the right Y-axis indicates the relative percentage of 6mA (based on the total number of adenine bases present in the assembled genome). Data represent the mean ± SEM of three biological replicates. ***(B)*** Pearson Correlation Coefficient heatmap showing the similarity between SMRT sequencing replicates. ***(C)*** Venn diagrams were used to depict the overlap in the profiles of 6mA across different timepoints, after merging replicates. ***(D)*** A Circos plot was used to visualize the genome-wide distribution of 6mA. The outermost black bars represent the 14 chromosomes. Blue and red bars represent genes on the positive and negative strands, respectively. The black trace represents the GC-richness profile. ***(E)*** Motif enrichment analysis of all 6mA-modified regions of the *P. falciparum* genome across all timepoints using weblogo v3. ***(F)*** Di-nucleotide context of 6mA in the *P. falciparum* genome across all timepoints.

Overlap analysis across the four timepoints revealed that 34.4% of 6mA loci are stably maintained throughout *P. falciparum* IED, 17.06% in three stages and 18.62% in two stages, with 5.4%, 5.5%, 6.1%, and 12.3% of the sites being unique to 10, 20, 30 and 40 hpi, respectively (**Fig. 2C**). The stable nature of 6mA during the parasite asexual developmental cycle is also apparent in the Circos plot of **Fig. 2D**; for instance, some 6mA peaks in chromosomes 12 and 14 (**Fig. 2D**; black arrowheads) are conserved across all four timepoints, while a 6mA signal in chromosome 14 (**Fig. 2D**; grey arrowheads) is specific only to the 10 and 20 hpi timepoints.

Lastly, we performed motif enrichment analysis using weblogo v3 [45] which revealed a consensus sequence of trAAA(A/T)G**<u>A</u>**(G/A/C)(T/G)(G/T)AA for 6mA in the *P. falciparum* genome (**Fig. 2E**). This was supported by an analysis of the dinucleotide context of 6mA, which identified GpA dinucleotides as the most frequently modified, followed by ApA, ApG and ApC (**Fig. 2F**). Of note, this pattern is more similar to *C. elegans* and humans, wherein GA-rich motifs (GAGG/AGAA and [G/C]AGG[C/T], respectively) are the primary 6mA hotspots [22], [27], and is in contrast to other unicellular eukaryotes where 6mA occurs predominantly at palindromic ApT dinucleotides [20], [46].

### 6mA is enriched in coding regions of the *P. falciparum* genome

We, thereafter, examined the genomic features associated with 6mA at the four IED timepoints, and observed that over 50% of the 6mA loci in the *P. falciparum* genome are located within exons of protein-coding genes, followed by approximately 20% within intergenic regions, nearly 17% in 5’ and 3’ UTRs, and the remaining 13% of loci distributed across intronic regions, ncRNA genes, pseudogenes, and other genomic features. (**Fig. 3A**). The exonic enrichment of 6mA prompted us to investigate the protein-coding genes that are marked with this DNA base modification, leading to the identification of 2,167 6mA-containing genes at 10 hpi, 1,867 at 20 hpi, 1,977 at 30 hpi, and 2,406 at 40 hpi (**Fig. 3B**). Amongst these, 942 genes were consistently methylated across all four timepoints (**Fig. 3C**) and encode for proteins such as AP2 domain transcription factors, ATP-dependent RNA helicases, and DNA repair proteins. (**Supplementary Table S3**). A metagene analysis of the average 6mA density profile across the coding sequence (CDS) of modified core genes further substantiated hypermethylation of exonic regions (**Fig. 3D**), with the non-template strand possessing more 6mA modifications as compared to the template strand across all timepoints (**Supplementary Fig. S5**).

**FIGURE 3.**
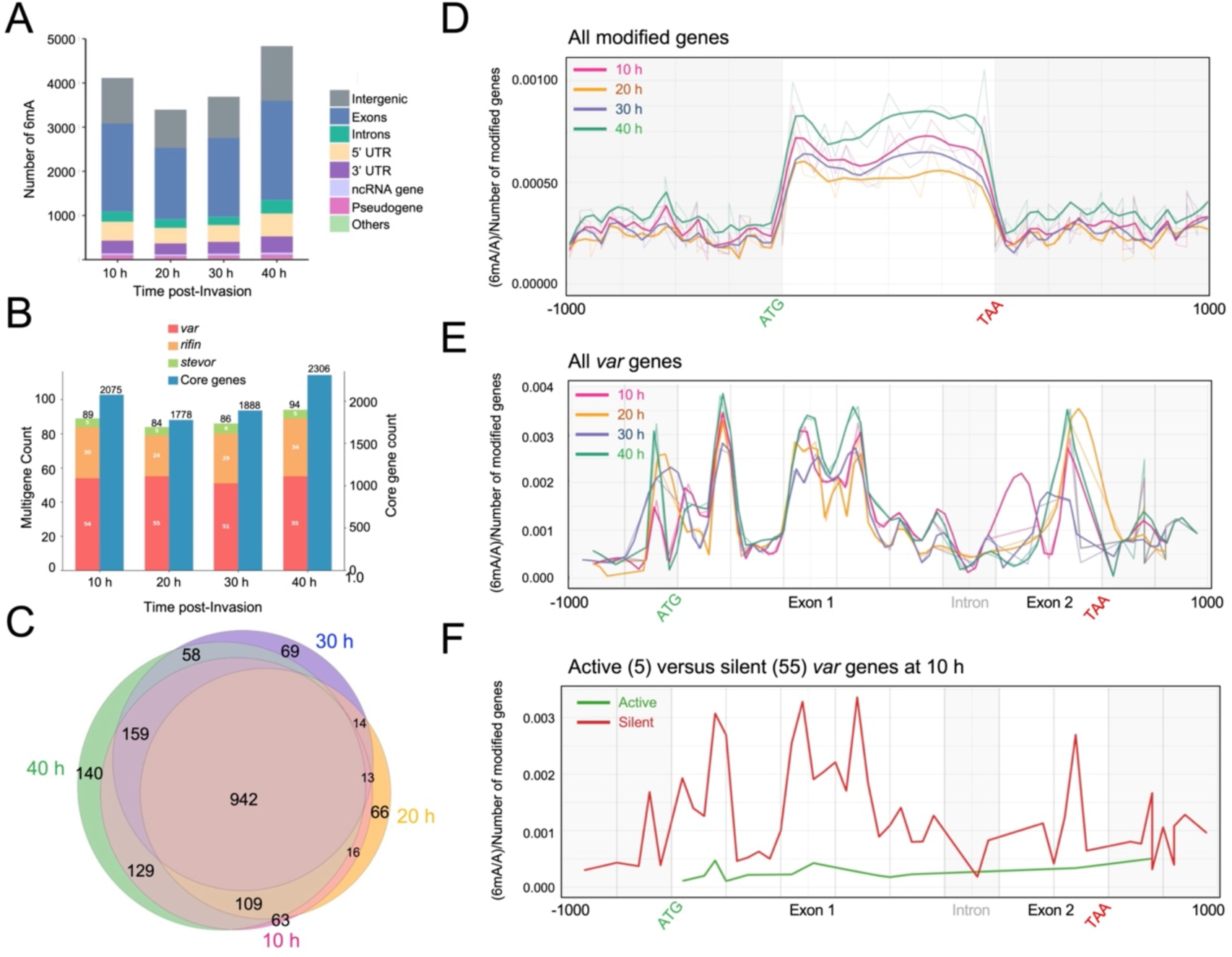
6mA is enriched in exonic regions of *P. falciparum* protein-coding genes and in exon 1 of silent *var* genes. ***(A)*** Stacked bar plots showing the distribution of 6mA modifications across different genomic features at four timepoints of *P. falciparum* IE development. UTR = Untranslated region; ncRNA = noncoding RNA. Exons (blue) and intergenic regions (grey) dominate. ***(B)*** Bar plots showing the number of genes containing 6mA at each timepoint. The left Y-axis represents the count of modified *var, rifin* and *stevor* genes while the right Y-axis represents the count of *P. falciparum* protein-coding genes that do not belong to these multigene families. ***(C)*** Venn diagrams were used to depict the overlap in the genes containing 6mA across different IE timepoints. ***(D)*** 6mA levels across the start codon (ATG), gene body, stop codon (TAA) and 1000 bp upstream and downstream flanks of modified genes that do not belong to multigene families. Each region was divided into 20 equal-sized bins for standardized representation. ***(E)*** 6mA levels across exon 1, intron, exon 2 and 1000 bp upstream and downstream flanks of all *var* genes at the four timepoints analysed in this study. ***(F)*** 6mA levels across exon 1, intron, exon 2 and 1000 bp upstream and downstream flanks of 55 silent *var* genes (red) and 4 active *var* genes (green) at 10 h. For ***E*** and ***F***, each genomic region was split into the appropriate number of bins based on relative sizes.

Notably, select members of multigene families such as *var*, *rifin* and *stevor* which encode for virulence surface antigens and are known to be epigenetically regulated [47] are marked with 6mA at all four timepoints (**Fig. 3B**). Analysis of the average 6mA distribution profile across the CDS and 1000 bp flanking regions of these genes demonstrated a genic enrichment profile (**Supplementary Fig. S6**), similar to the core genes (**Fig. 3D**). In the specific case of the 60-member *var* gene family, the central portions of exons 1 and 2 were observed to be hypermethylated as was the 5’UTR, with the intron being hypomethylated (**Fig. 3E**). Moreover, a comparison of active (5 members including *PF3D7_0412700, PF3D7_0632800, PF3D7_1240600, PF3D7_0632500,* and *PF3D7_0412400*) and silent *var* genes (the remaining 55 members), which were categorised based on RNA-seq analysis of the three 10 hpi biological replicates used for SMRT sequencing (**Supplementary Fig. S7** and **Supplementary Table S4**), revealed a higher prevalence of 6mA modifications in silent *var* genes at 10 hpi, with active genes containing negligible levels of 6mA within the CDS or the 1000 bp upstream region that corresponds to the 5’UTR (**Fig. 3F**).

### 6mA-containing *P. falciparum* genomic regions do not occlude nucleosomes

Studies in ciliates such as *T. thermophila* and *O. trifallax* have demonstrated that 6mA preferentially localizes to nucleosome-depleted linker DNA, disfavours nucleosome occupancy and promotes an open chromatin state [21], [30], [48]. To investigate whether a similar regulatory relationship exists in *P. falciparum*, processed MNase-seq data for the *P. falciparum* 3D7 strain at 10, 20, 30 and 40 hpi were obtained from publicly available databases [49], [50] and compared to 6mA positions derived from SMRT sequencing data. Contrary to other ancient eukaryotes, our analysis indicated that 6mA-containing regions of *P. falciparum* chromosomes are enriched in nucleosomes (**Fig. 4A** and **Supplementary Fig. S8A**; red line). This was further confirmed by analysing the distribution of nucleosomes across adenines randomly sampled across the genome (**Fig. 4A** and **Supplementary Fig. S8A**; black line), and the lower levels of 6mA found within *P. falciparum* genomic regions that correspond to linker DNA (**Fig. 4B** and **Supplementary Fig. S8B**).

**FIGURE 4.**
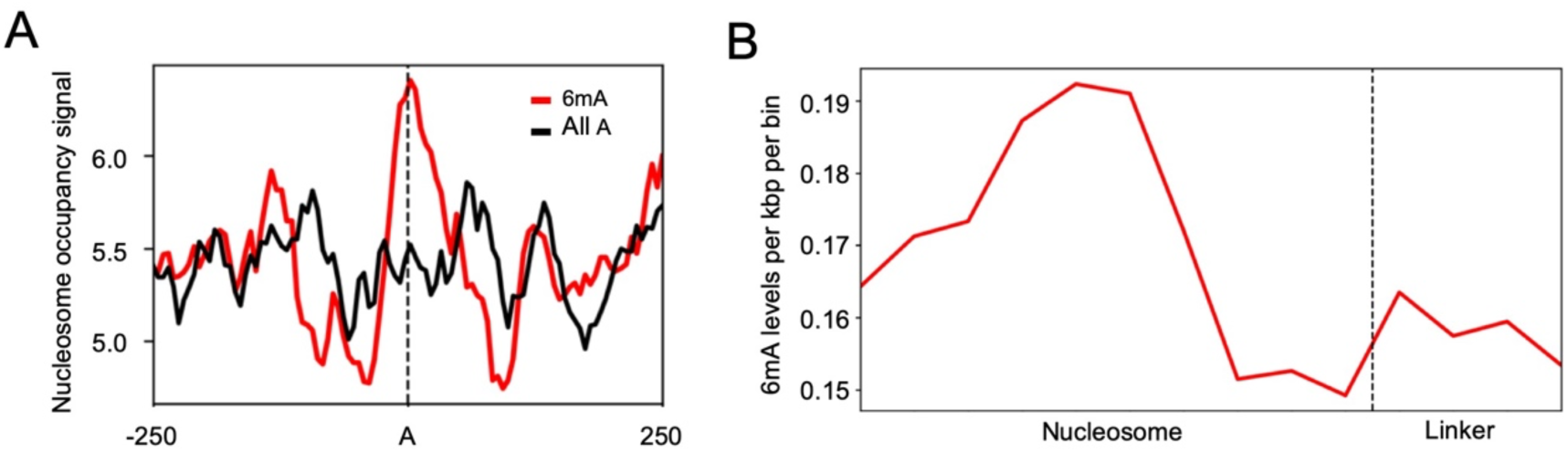
6mA loci in the P. falciparum genome do not occlude nucleosomes. **(A)** The X-axis represents an adenine base anchored at “0” and its +/- 250 bp flanking regions. The Y-axis region represents the nucleosome occupancy signal obtained from Kensche *et al.,* 2016 [49]. The red line is centred at 6mA loci while the black line is centred at an equal number of adenine sites chosen at random. **(B)** The X-axis represents the position of the nucleosome (divided into 10 bins) and linker (divided into 4 bins). The Y-axis represents 6mA levels per kbp per bin.

### 6mA may activate gene expression during *P. falciparum* IED

Ultimately, to link adenine methylation to epigenetic regulation of gene expression during *P. falciparum* IED, the distribution patterns around 6mA loci of six histone PTMs that are associated with active transcription, including histone H3 lysine 4 dimethyl (*i.e.,* H3K4me2), H3K4me3, H3K9ac, H3K14ac, H3K27ac, and H4ac, and four that are linked to transcriptional silencing, primarily of multigene families, namely H3K9me3, H3K27me3, H3K36me2, and H3K36me3, were assessed at 20 and 40 hpi. The profiles of the activating and silencing histone marks were re-calculated from published ChIP-seq datasets for the *P. falciparum* 3D7 strain [51], [52], [53], with the analysis focusing on 6mA sites located within the 5′UTR and CDS (inclusive of exonic and intronic regions) of core genes. As shown in **Fig. 5A** and **Supplementary Fig. S9A** for 20 hpi and 40 hpi, respectively, H3K4me2 and H3K4me3 signals are enriched approximately 100 bp downstream of the 6mA locus within 5′UTRs, while, within CDSs, a strong correlation is evident between 6mA positions and activating histone marks, particularly H3K9ac and H3K14ac. Conversely, silencing histone PTMs are not enriched around 6mA positions in either 5’UTRs or CDSs (**Fig. 5B** and **Supplementary Fig. S9B**); in fact, H3K27me3 appears to be depleted from nucleosomes in genomic regions marked with 6mA.

**FIGURE 5.**
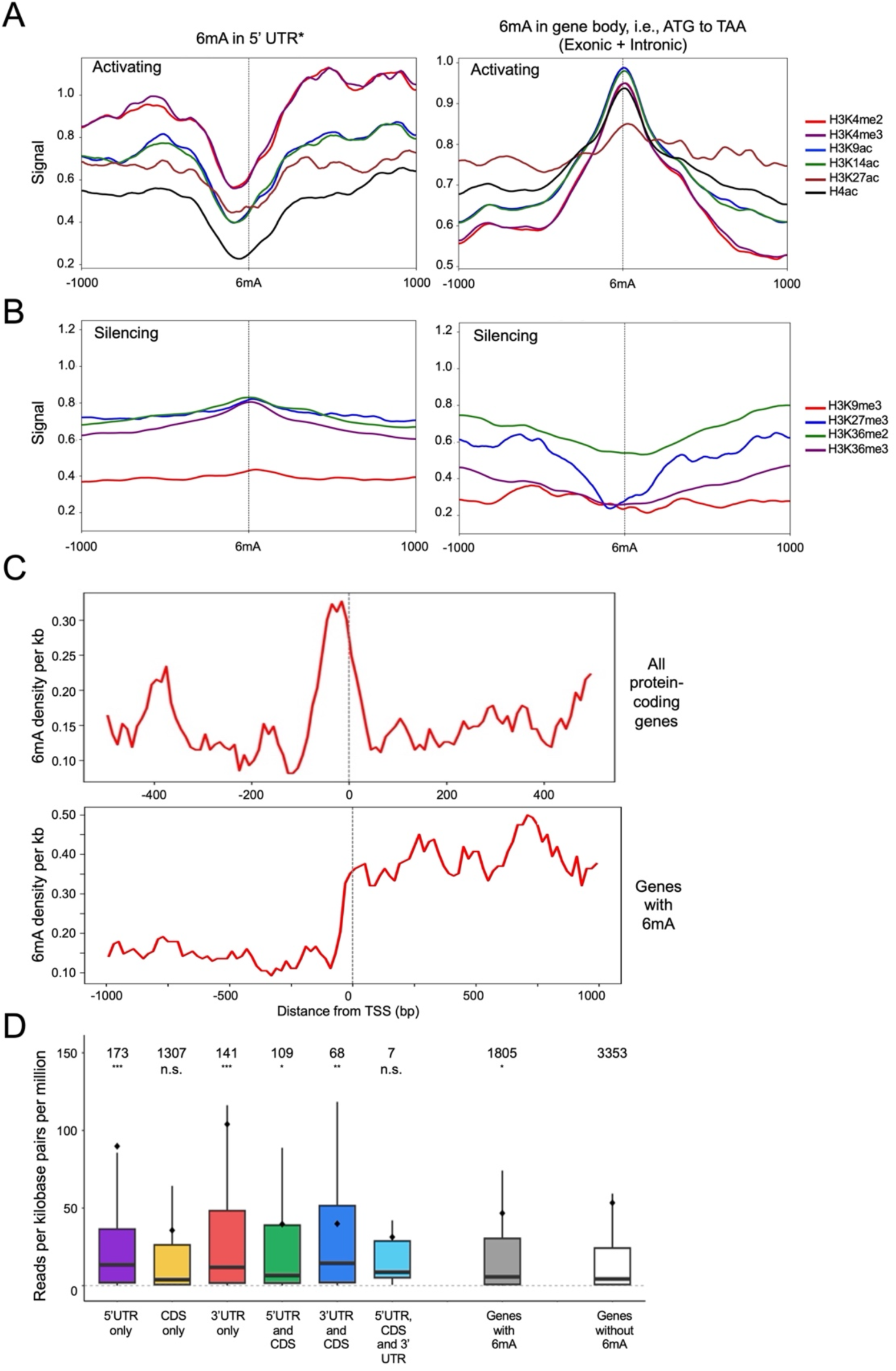
Genic 6mA loci correlate with active histone epigenetic marks and increased steady state transcript levels during *P. falciparum* asexual development. ***(A & B)*** DeepTools plotprofile was used to compare the read density distribution of the indicated activating ***(A)*** and silencing **(B)** histone PTMs[51], [52] for all 6mA positions located within 5’UTRs (*left*) or gene bodies (*right*) of modified genes at 20 h. ***(C)*** DeepTools was used to analyse the density of 6mA +/-500 bp from Transcription Start Sites (TSS; https://plasmodb.org) for 20 h data. *Top panel:* All TSS; *Bottom panel:* TSS of modified genes alone. ***(D)*** Box plots were used to represent the distribution of normalised RNA-seq read counts (Reads per kilobase per million mapped reads (RPKM)) for 6mA-modified genes at 20 h. RNA-seq data were obtained from Kensche *et al.,* 2016 [49]. The box shows the interquartile range, with the line inside representing the median; the mean is indicated using a ♦. The numbers of genes in each group are shown. p-values were calculated using a Wilcoxon test. UTR = Untranslated region; CDS = Coding sequence.

The observation that 6mA may be preferentially associated with transcriptionally active chromatin encouraged us to analyse 6mA density around transcription start sites (TSS) for all 5300 *P. falciparum* protein-coding genes for all timepoints, which revealed that, on average, 6mA signals are enriched at positions-50 to-10 upstream of the TSS (**Fig. 5C** and **Supplementary Fig. S10;** upper panel). In the specific case of 6mA-modified genes, *i.e.,* genes with 6mA in 5’UTR, CDS and/or 3’UTR, 6mA density starts increasing approximately 10 bp upstream of the TSS and stays elevated in the 500 bp downstream flank (**Fig. 5C** and **Supplementary Fig. S10;** lower panel). This, in turn, results in significantly higher steady state transcript levels of 6mA-modified genes relative to unmodified genes, especially when the modifications are in the 5’ or 3’ UTRs (**Fig. 5D** and **Supplementary Fig. S11**); for this comparative analysis, RNA-seq-derived transcriptomic data from a published study [49] was used. Overall, during *P. falciparum* IED, our observations point to a transcriptional activating role for adenine DNA methylation for core parasite genes.

## DISCUSSION

The role of 6mA in genome regulation has been extensively characterized in prokaryotes, where it participates in processes such as restriction–modification systems, DNA mismatch repair, and transcriptional regulation [54], [55]. In contrast, its function in eukaryotes has only recently emerged and remains context-dependent, with evidence ranging from gene activation and nucleosome positioning in unicellular organisms to the ongoing debate regarding its prevalence and functional relevance in more recently evolved eukaryotes [20], [22], [25], [30], [43], [56]. Here, we provide multiple lines of evidence supporting the presence and potential regulatory role of 6mA in the AT-rich genome of the malaria parasite *P. falciparum*, thereby positioning 6mA as an evolutionarily conserved, yet mechanistically diverse, epigenetic mark.

A major strength of our study lies in the orthogonal validation of 6mA using complementary methodologies, including antibody-based detection, LC-MS/MS quantification, and SMRT sequencing. Indeed, while the concordance between LC-MS/MS and SMRT sequencing estimates supports a relatively low abundance of 6mA (approximately 0.02 to 0.05%) in the *P. falciparum* genome, placing it within the lower range reported for eukaryotes but comparable to earlier observations in this parasite [41], the convergence of the three approaches alleviates concerns raised in previous studies about false positives arising from antibody cross-reactivity or sequencing artefacts [14], [41], [57]. Furthermore, the observed enrichment of 6mA within an A-rich consensus sequence (AAA(A/T)GA(G/A/C)(T/G)(G/T)AA) in *P. falciparum*, with a notable preference for GpA dinucleotides, is in contrast to other unicellular eukaryotes where 6mA is enriched at ApT-rich motifs [20], [21], [30], [46]. Indeed, the *P. falciparum* motif is more extended and degenerate, aligning closely with the GA-centric motifs of metazoans [22], [23]. This may reflect species-specific targeting of the 6mA methyltransferase machinery or local structural features of *P. falciparum* DNA that favour methylation. Notably, unlike the well-defined palindromic motifs seen in prokaryotic restriction-modification systems [58], the motif identified here is less constrained, suggesting a more flexible or context-dependent targeting mechanism in *P. falciparum*. Further investigation will be required to determine whether this motif is directly recognized by PfN6AMT or arises as a consequence of chromatin accessibility and sequence composition.

One of the major findings of this study is the apparent stability of 6mA levels across *P. falciparum* IED. Unlike dynamic epigenetic marks such as histone PTMs, which undergo stage-specific reprogramming in *P. falciparum* [51], [59], [60], [61], 6mA appears largely invariant both in global abundance and genomic distribution. This suggests that, rather than acting as a transient regulatory signal, 6mA may function as a constitutive epigenetic feature associated with a subset of genes. A similar stability has been reported in certain unicellular eukaryotes, where 6mA marks persist across cell cycles and are linked to housekeeping or consistently expressed genes [20], [30]. In this context, 6mA in *P. falciparum* may contribute to maintaining basal transcriptional competence rather than driving stage-specific transcriptional changes. Consistent with this interpretation, our genome-wide analyses reveal a preferential enrichment of 6mA within gene bodies, particularly exons and UTRs, and a positive correlation with steady-state transcript levels. This association between genic 6mA and transcriptional activity mirrors observations in other eukaryotic systems, including *C. reinhardtii* and *T. thermophila*, where 6mA is enriched near TSSs and marks actively transcribed genes [20], [30]. Importantly, in *P. falciparum*, the enrichment of 6mA upstream of the TSS and within 5′UTRs suggests a potential role in transcriptional initiation or early elongation. The observed co-localization with activating histone marks such as H3K4me3 and H3K9ac further supports a model in which 6mA is integrated into the broader epigenetic landscape that defines transcriptionally permissive chromatin states [62] and mirrors patterns of 6mA-H3K4me3 co-occurrence that have been recently reported for other protists [46].

However, the relationship between 6mA and chromatin architecture in *P. falciparum* appears to diverge from that described in ciliates and other alveolates. In organisms such as *O. trifallax* and *T. thermophila*, 6mA is strongly associated with nucleosome-depleted linker DNA and is thought to promote nucleosome positioning by excluding histones [16], [21], [48]. In contrast, our data indicate that 6mA in *P. falciparum* is enriched within nucleosome-occupied regions, suggesting a fundamentally different mode of interaction with chromatin. This unexpected finding may reflect the unique genomic architecture of *P. falciparum*, which is characterized by extreme AT richness (∼80%) and atypical nucleosome organization [49]. It is plausible that, in this context, 6mA contributes to stabilizing nucleosomes or modulating nucleosome dynamics in a manner that supports transcription rather than opposing chromatin compaction. Such a model would represent a significant departure from the canonical view of 6mA as a marker of open chromatin and highlights the evolutionary plasticity of epigenetic mechanisms.

A particularly interesting observation from our study is the preferential enrichment of 6mA within silent *var* genes, in contrast to their actively transcribed counterparts, which exhibit minimal or negligible levels of this modification. The *var* gene family, which encodes the *P. falciparum* Erythrocyte Membrane Protein 1 virulence protein, is subject to mutually exclusive expression and tightly regulated epigenetic control, primarily mediated by histone modifications such as H3K9me3 and heterochromatin protein 1 [47], [63], [64]. The enrichment of 6mA in silent *var* loci suggests that, unlike its association with transcriptional activation in core genes, 6mA may play a context-dependent role within multigene families, potentially contributing to the maintenance of transcriptionally repressed states. One possibility is that 6mA participates in reinforcing epigenetic memory at silent *var* loci, thereby stabilizing their inactive state across replication cycles. Alternatively, 6mA may mark loci poised for activation, analogous to bivalent chromatin states described in other eukaryotic systems [20], [46], [65], although such a mechanism remains to be demonstrated in *P. falciparum*. Notably, the apparent divergence in 6mA function between core genes and *var* genes highlights the complexity of epigenetic regulation in the parasite and suggests that the functional outcome of 6mA deposition is likely shaped by local chromatin context and interacting regulatory factors. Further experimental interrogation, particularly through locus-specific analyses and perturbation of 6mA deposition, will be required to delineate whether 6mA actively contributes to *var* gene silencing or serves as a secondary consequence of heterochromatin formation.

Lastly, the identification and biochemical characterization of a putative DNA 6mA methyltransferase, PfN6AMT, which is essential for parasite IED [66], further strengthens the argument for a functional 6mA regulatory system in *P. falciparum*. PfN6AMT shares sequence and structural similarity with members of the METTL4 family, which have been implicated in DNA adenine methylation in diverse eukaryotes [27], [31]. The conservation of the catalytic NPPY motif and the loss of activity upon its mutation provide additional evidence that PfN6AMT functions as a *bona fide* methyltransferase. Yet, in light of a recent study of DNA 6mA pathways in 18 unicellular eukaryotes which established that genomic 6mA is restricted to protists that encode the AMT1 clade of MT-A70 methyltransferases [46] the apparent absence of a canonical AMT1 enzyme in *P. falciparum* raises the intriguing possibility that apicomplexan parasites have evolved a distinct enzymatic pathway for DNA adenine methylation. In this context and given the sequence similarity of PfN6AMT to hN6AMT1, the former may have been acquired from its host during the emergence of parasitism and may regulate a divergent *P. falciparum* 6mA pathway. Nonetheless, the debate about hN6AMT1’s DNA methyltransferase activity [27], [32], [33], [67] warrants a detailed genetic and biochemical study to establish the “true” contribution of PfN6AMT to genomic 6mA deposition in *P. falciparum in vivo*. Moreover, while *P. falciparum* encodes an MT-A70 orthologue PfMT-A70, this protein is METTL3-like and functions as an mRNA m^6^A methyltransferase [68], and its role in DNA 6mA deposition has not been explored. Indeed, it is possible that PfN6AMT and PfMT-A70 work together for *de novo* versus maintenance DNA adenine methylation during *P. falciparum* growth and replication, as has been reported for *Tetrahymena* by AMT2/AMT5 and AMT1, respectively [69], [70].

We recognize that our results supporting an epigenetic role for 6mA remain correlative, not causal. Genetic perturbation of PfN6AMT (and PfMTA-70), coupled with 6mA, transcriptomic and chromatin profiling, will therefore be essential to determine whether this DNA base modification directly regulates gene expression or merely marks transcriptionally permissive loci; this work is ongoing. Additionally, the relatively low abundance of 6mA in *P. falciparum*, despite its genomic AT-richness, raises the possibility that even subtle technical biases could influence detection, underscoring the importance of continued methodological refinement. In this regard, our use of a stringent 25X coverage threshold for 6mA calling from our SMRT sequencing data, while prioritizing high-confidence 6mA loci, may have excluded low-frequency or heterogeneous 6mA events that occur in only a subset of parasite populations or at specific developmental states. Future studies employing increased sequencing depth, improved base modification detection algorithms, and single-cell or single-molecule approaches may provide a more complete view of the dynamic and potentially heterogeneous 6mA landscape in *P. falciparum*. Finally, the interplay between 6mA and other epigenetic layers, including RNA modifications such as m6A, histone post-translational modifications, and histone variants, remains unexplored and represents an important avenue for future research.

In conclusion, our study provides substantial, orthogonal evidence that 6mA is a functionally relevant DNA base modification in *P. falciparum*, albeit of low abundance, and associated with transcriptionally active genes. The distinct chromatin associations of 6mA in this pathogen, *i.e.,* its presence in genomic regions that are occupied by nucleosomes that contain transcriptionally permissive histone PTMs, underscore the evolutionary diversity of epigenetic regulation in closely related protists. Ultimately, the elucidation of the precise molecular functions of 6mA and its associated enzymatic machinery in *P. falciparum* may not only open new avenues for investigating gene control mechanisms in malaria parasites, it may also reveal novel targets for therapeutic intervention.

## MATERIALS AND METHODS

### Parasite culture and genomic DNA isolation

Asexual blood stages of the *P. falciparum* 3D7 laboratory strain were grown and cultured as previously described [71], [72]. For south-western dot blot analysis, genomic DNA was isolated using the DNeasy Blood and Tissue kit (Qiagen) with RNase A treatment as per manufacturer’s instructions. For mass spectrometry and SMRT seq, the MagAttract HMW DNA kit (Qiagen) was used with RNase A treatment during isolation. Briefly, packed blood was subjected to a series of four RPMI-incomplete washes to generate WBC-free growth conditions. Next, parasite cultures were initiated in the washed blood using a frozen inoculum of synchronized ring stages. To tighten the growth window, D-sorbitol (Sigma-Aldrich) was used, which synchronizes parasite cultures at the ring stage. Next, Percoll (Sigma-Aldrich) treatment was carried out to enrich segmented schizonts, following which the culture was monitored every two hours. Samples were accordingly harvested at 10, 20, 30 and 40 hpi and subjected to saponin treatment to release free parasites, which were lysed according to the MagAttract HMW DNA kit protocol and subjected to genomic DNA extraction. The quality of the genomic DNA, particularly the presence of HMW fragments, was assessed by agarose gel electrophoresis while DNA concentration was measured using Thermo Scientific NanoDrop One/One^C^.

### Preparation of nuclear extracts

Parasite nuclear components were extracted as described by [72]. 5 × 10^9^ schizont stage parasites were extracted from infected erythrocytes using saponin lysis, resuspended in 1 ml of lysis buffer (10 mM HEPES pH 7.9, 10 mM KCl, 0.1 mM EDTA, 0.1 mM EGTA, 1 mM DTT, 0.65% NP-40) supplemented with protease inhibitors (G-BIOSCIENCES) and incubated at 4°C for 30 minutes. Complete parasite lysis was accomplished with 200 strokes using a pre-chilled Dounce homogenizer. The lysate was centrifuged at 14,000 rpm for 10 minutes at 4°C after which the supernatant, containing the cytoplasmic fraction, was discarded. The resulting pellet was subjected to three washes with phosphate-buffered saline (PBS) and subsequently resuspended in 100 ml of extraction buffer (20 mM HEPES pH 7.9, 400 mM NaCl, 1 mM EDTA, 1 mM EGTA, 1 mM DTT) supplemented with protease inhibitors, followed by incubation with vigorous shaking for 30 minutes at 4°C. Finally, after centrifugation at 14000 rpm for 10 minutes at 4°C, the supernatant corresponding to the nuclear extract was collected. The concentration of the extracts was determined using a Bradford assay.

### South-western blotting with anti-6mA antibodies

Genomic DNA was denatured at 95°C and immediately transferred to ice to prevent renaturation. DNA concentrations ranging from 400 to 25 ng were spotted onto a positively charged nylon membrane (HIMEDIA) and UV cross-linked at 70000 μJ/cm² for one minute, followed by blocking using 5% milk in PBS + 0.1% Tween-20 (PBST). The membrane was probed using rabbit monoclonal antibodies against 6mA (Synaptic Systems) at 1:5000 dilution overnight at 4°C. The next day, the membrane was incubated with horseradish peroxidase-conjugated anti-rabbit secondary antibodies at 1:5000 dilution in PBST for one hour at room temperature. The ECL Western Blotting Detection Reagent Clarity Kit (Bio-Rad 170-5060) was used for signal detection. Images were captured using BIO-RAD ChemiDoc^TM^ MP Imaging System.

### Plasmid constructs and recombinant protein purification

The coding sequence of the putative methyltransferase (*PF3D7_1303100*) was PCR-amplified from *P. falciparum* genomic DNA and cloned into the bacterial expression vector pET28a in-frame with an N-terminal hexahistidine (6XHis) tag while the coding sequence of human N6AMT1 (UniProt ID: Q9Y2K6) was PCR-amplified from human cDNA and cloned into the bacterial expression vector pET22b, in-frame with C-terminal 6XHis tag (**Supplementary Table S5**). A catalytically dead version of *PF3D7_1303100* was created using site-directed mutagenesis. Briefly, forward and reverse primers containing the desired mutation were designed (**Supplementary Table S5**) and used to amplify two fragments of the *PF3D7_1303100* gene, which were then joined using SOE-PCR and cloned into pET28a. Diagnostic restriction enzyme digestion and Sanger sequencing were performed to confirm all plasmid clones.

*Escherichia coli* BL21 DE3 cells were used to express recombinant proteins. Protein induction was carried out using 0.3 mM IPTG for 8 h at 18°C, with the induced 6XHis-tagged proteins purified using Ni-NTA Agarose (Qiagen) affinity chromatography. Since 6XHis-PfN6AMT and the mutant were insoluble, they were purified using a denaturation refolding protocol. Briefly, cell pellets were resuspended in Buffer A (100 mM monosodium phosphate, 10 mM Tris-HCl, 6 M guanidinium hydrochloride, pH 8.0) at 5 ml per gram wet weight and stirred for 2 hours at room temperature to solubilize inclusion bodies. The lysate was clarified by centrifugation at 10,000 × g for 30 minutes at 4°C, and the supernatant was collected. The clarified lysate was loaded onto a pre-equilibrated Ni-NTA column under gravity flow to allow binding of the 6XHis-tagged protein under denaturing conditions. The column was then washed three times each with Buffer A and Buffer B (100 mM monosodium phosphate, 10 mM Tris-HCl, 8 M urea, pH 8.0) to remove non-specifically bound proteins. On-column refolding was achieved by sequentially washing the column with a stepwise decreasing urea gradient using seven refolding wash buffers (all containing 20 mM Tris-HCl, 20 mM imidazole, 1 mM β-mercaptoethanol, pH 8.0) supplemented with 6 M, 5 M, 4 M, 3 M, 2 M, 1 M, and 0 M urea, respectively (Refolding Wash Buffers 1–7), allowing the immobilized protein to gradually adopt its native conformation. The refolded protein was subsequently eluted using a stepwise imidazole gradient (Elution Buffers 1–4; all containing 20 mM Tris-HCl, 500 mM NaCl, 1 mM β-mercaptoethanol, pH 8.0) with imidazole concentrations of 100 mM, 200 mM, 300 mM, and 400 mM, respectively, and fractions were collected.

6XHis-hN6AMT1 was purified from soluble fractions as follows: cell pellets were resuspended in lysis buffer (100 mM Tris-HCl, 150 mM NaCl, 1% Triton X-100, 5% glycerol, 10 mM imidazole, pH 8.0) and lysed by sonication (Qsonica Q500). The clarified cell lysate was obtained by centrifugation and loaded onto a pre-equilibrated Ni-NTA column under gravity flow. The column was washed sequentially with three wash buffers to remove non-specifically bound proteins: Wash Buffer 1 (100 mM Tris-HCl, 150 mM NaCl, 1% Triton X-100, 5% glycerol, 10 mM imidazole, pH 8.0), Wash Buffer 2 (100 mM Tris-HCl, 150 mM NaCl, 5% glycerol, 10 mM imidazole, pH 8.0), and Wash Buffer 3 (100 mM Tris-HCl, 500 mM NaCl, 5% glycerol, 10 mM imidazole, pH 8.0). The target protein was eluted using a stepwise imidazole gradient (100–400 mM imidazole in 100 mM Tris-HCl, 150 mM NaCl, 5% glycerol, pH 8.0), and fractions were collected.

Following elution, fractions containing the target protein were pooled and subjected to dialysis using a Sigma Pur-A-Lyzer dialysis tube. For 6XHis-PfN6AMT or its mutant, on-column refolding was further continued during dialysis by applying a stepwise decreasing urea gradient. The protein was dialyzed against 2 liters of dialysis buffer (10 mM Tris-HCl, 50 mM NaCl, 5 mM β-mercaptoethanol, 2% glycerol, pH 7.8) at 4°C with buffer replacement every 2 hours using sequentially decreasing urea concentrations: 2 M, 1.5 M, 1 M, 0.5 M, and 0 M urea, respectively, over a total of 10 hours. For 6XHis-hN6AMT1, the protein was dialyzed against 2 liters of dialysis buffer at 4°C for 8 hours with buffer replacement every 2 hours to ensure complete removal of imidazole. Post-dialysis, protein purity and concentration were assessed by Coomassie Brilliant Blue staining and Bradford assay, respectively.

### Methyltransferase activity assay

Methyltransferase activity assays were performed as previously described with minor modifications [27]. Briefly, the reactions were carried out in a 50 µl reaction volume containing 1 µg of recombinant protein or 10-20 µg of nuclear extracts together with 250 pmol of DNA oligonucleotides (**Supplementary Table S5**), 800 mM S-adenosylmethionine, 50 mM Tris-HCl (pH 7.6), 50 mM KCl, 10 mM Mg(OAc), 100 mg/ml bovine serum albumin and 2.7 mM β-mercaptoethanol. Reactions were incubated overnight at 25°C, followed by protein inactivation at 95°C for 10 minutes after which the DNA was cleaned up using the Nucleospin® Macherey-Nagel PCR purification kit, quantified and analysed using anti-6mA dot blotting.

### Phylogenetic analysis

The amino acid sequences of N6AMTs (UniProt ID: Q8IET4 and Q9Y2K6) were obtained from UniProt [73] while the other protein sequences used for phylogenetic tree construction were acquired from a previous study (Table S2 of Beh et al., 2019 [21]), with tree construction performed as described [21].

### Quantification of 6mA in genomic DNA by UHPLC-QQQ-MS/MS

Genomic DNA (1000 ng per sample) was enzymatically digested to nucleosides using Nucleoside Digestion Mix (New England Biolabs, M069S), a blend of enzymes that breaks DNA into its individual nucleosides, at 37 °C for 2 hours [42]. Following digestion, both samples and standards [2’-deoxyadenine (dA), the unmodified DNA nucleoside, and N⁶-methyl-2’-deoxyadenine (6mA), its methylated form] were diluted to 100 µL with double-distilled water, then filtered through 0.22 µm syringe filters. Finally, 5 µL of each filtrate was injected for analysis. Separation was performed on an Agilent 1290 UHPLC (ultra-high performance liquid chromatography) system equipped with a C18 reversed-phase column (2.1 × 50 mm, 1.8 µm). Mobile phase A was water with 0.1% (v/v) formic acid, and mobile phase B was methanol with 0.1% (v/v) formic acid. Mass spectrometry was performed using an Agilent 6470 triple quadrupole (QQQ) mass spectrometer in positive electrospray ionisation (ESI) mode with dynamic multiple reaction monitoring (dMRM), a method for detecting specific molecules. Transitions monitored were m/z (mass-to-charge ratio) 252.1 → 136.0 for dA and 266.1 → 150.0 for 6mA. 6mA/dA ratios were determined from calibration curves prepared by serially diluting N6-methyladenine (6mA) and deoxyadenine (dA) standards. To account for background, a negative control (“mock” digestion) containing all reagents, but no DNA was used to measure background levels of 6mA and dA; these values were then subtracted from sample measurements.

### SMRTbell library preparation, SMRT sequencing and SMRT data processing

A total of 12 genomic DNA samples, including three biological replicates each of four *P. falciparum* IED timepoints, 10, 20, 30 and 40 hpi, were multiplexed and analyzed by SMRT sequencing. 1 mg of high molecular weight genomic DNA sheared to ∼15 kbp fragments using g-tubes (Covaris) was used as input for SMRTbell template preparation. Quality and quantity of the input was determined using an Agilent Tape Station 4150 with a genomic DNA kit (Agilent Technologies) and Qubit 1X high-sensitivity double-stranded (ds) DNA solution (ThermoFisher), respectively. SMRTbell library preparation was performed according to the specifications from the manufacturer using the Express Template Kit 2.0 (Pacific Biosciences). Briefly, the sheared genomic DNA underwent DNA damage repair to repair any potential nicked DNA, followed by an End Repair reaction to fill in any single stranded ends and add a A-tail to enable the next blunt-end ligation step. This repaired material was then ligated to symmetrical barcoded adapters on both ends of the molecule to facilitate multiplexing libraries downstream. Following adapter ligation, the material was then subjected to digestion with a nuclease mix, allowing for digestion of unligated genomic DNA, leaving only adapter bound material for sequencing. This final library was assessed for quantity and quality using the TapeStation genomic DNA and Qubit 1X HS dsDNA assays, as described above. Size and concentration values were used to generate an equimolar pool of all 12 barcoded samples prior to sequencing. Once pooled, the SMRTbell libraries were annealed to sequencing primer, which binds to the single-stranded portion of each barcoded adapter. Following annealing, they were bound to the Revio sequencing polymerase (v1) and then underwent a magnetic-bead based wash to remove any unbound complex or excess polymerase. Final size and concentration of the multiplexed libraries were input into the Sample Preparation module of the SMRTLink web portal to obtain proper calculations for the final loading complex. The polymerase bound complex was loaded and run on one Revio SMRT cell using a 30-hour movie with polymerase kinetics metrics collected alongside sequencing data.

Following data collection, circular consensus (CCS) sequencing reads were generated on the instrument by collapsing multiple reads of the same molecule into one highly accurate intramolecular consensus sequence per molecule. Those reads that have an empirical sequence accuracy of >99% are categorized as “high fidelity” or “HiFi” reads and were used for all downstream analyses. All reads were exported from the instrument (**Supplementary Table S6**), and secondary analyses was performed using the SMRTLink (v12) SMRTAnalysis suite. Genome assembly was first performed using the Microbial Genome Analysis tool with default settings. Following this, the Base Modification and Find Base Motifs tools were run on the raw data using default settings, with the newly assembled genome serving as reference for each sample. For downstream analyses, the basemods.gff file was filtered to retain Base Modification calls with minimum 25X coverage.

## 6mA data analysis

### 6mA position extraction and mapping

The basemods.gff file provided 41-nucleotide sequence centered on each candidate modification site, along with chromosome, contig, strand and position information. These 41-nucleotide sequences were converted to FASTA format and aligned to the *P. falciparum* 3D7 reference genome (PlasmoDB release 68 genome.fasta; https://plasmodb.org) to generate Sequence Alignment Map (SAM) files, which were subsequently converted to Binary Alignment map (BAM) format, sorted and indexed using samtools [74]. The three replicate BAM files per timepoint were merged into a single BAM file by considering modified loci that were identified in at least two replicates and converted to BED format, using bedtools. Finally, duplicate genomic positions were removed from each merged BED file, retaining only one read per unique chromosomal coordinate, producing a non-redundant set of high-confidence modification sites for each timepoint for downstream analysis.

### Feature correlation

To characterize the genomic distribution of modified sites, high-confidence 6mA loci were annotated against known genomic features of the *P. falciparum* 3D7 genome (PlasmoDB-68_Pfalciparum3D7.gff; https://plasmodb.org). Each modification site was assigned to one of the following feature categories: exons, introns, 5’UTR, 3’UTR, ncRNA, pseudogenes and intergenic regions. The number of 6mA modifications per feature class was counted to visualize the feature-level distribution of 6mA sites across the IED cycle.

### Gene Body Enrichment Analysis

To examine the distribution of modified sites relative to gene structure, gene body enrichment analysis was performed in R using custom scripts alongside the tidyverse, furrr, and Biostrings packages [75]. CDS coordinates were extracted from PlasmoDB-68_Pfalciparum3D7.gff, spanning the full gene body from start codon (ATG) to stop codon (TAA). Each gene body was divided into 20 equal bins, flanked by 1,000 bp upstream and downstream regions also divided into 20 bins, yielding a standardised positional profile. Modification sites were intersected with each bin, with bin numbering adjusted for strand orientation to align all genes 5’ to 3’. Modification density per bin was normalised by the total count of the relevant unmodified base within that bin (derived from the reference genome using Biostrings) and divided by the total number of genes, yielding an average modification density per bin. Parallel processing via furrr was used to accelerate computation. Smoothed profiles were generated using LOESS regression and visualised with ggplot2 in R [76] separately for template and non-template strands. This analysis was performed separately for non-multigene families and *var* gene families. For *var* genes, a two-exon bin structure with 1,000 bp flanking regions was applied to reflect their distinct genomic architecture, and modification density was compared between transcriptionally active and silent *var* genes to assess differential 6mA deposition.

### Histone PTM correlation

To investigate the chromatin environment surrounding 6mA modification sites, publicly available ChIP-seq datasets for *P. falciparum* were obtained from GEO (GSE63369, GSE47349;[51], [53]). Raw ChIP-seq reads were quality assessed using FastQC and adapter-trimmed using Trim Galore [77] Trimmed reads were aligned to the *P. falciparum* 3D7 reference genome using BWA-MEM [78], and peaks were called using MACS2 [79]. Normalised bigWig coverage tracks were generated for each histone mark. Read density profiles of activating (H3K4me2, H3K9ac, H3K14ac, H3K4me3, H3K27ac, H4ac) and repressive (H3K9me3, H3K27me3, H3K36me2, H3K36me3) histone PTMs were plotted centred on individual 6mA sites using deepTools plotProfile [80]. Profiles were computed separately for 6mA sites located within 5’UTRs and within gene bodies (CDS; ATG to TAA, exonic and intronic), over a ±1,000 bp window divided into 200 bins, using a custom Python wrapper script. The X-axis represents the distance from each 6mA site and the Y-axis represents ChIP-seq signal strength of each PTM. Activating and repressive marks were plotted separately to distinguish chromatin states associated with 6mA-marked loci in transcriptionally active versus silent genes. A smoothening window of 7 bins was applied for visualization.

### Nucleosome occupancy signatures

To assess the relationship between 6mA modifications and nucleosome positioning, publicly available MNase-seq nucleosome occupancy data [49], [50] were compared against 6mA sites across all four timepoints (10 h, 20 h, 30 h, 40 h) using two custom Python scripts. The nucleosome occupancy signal was plotted centred on individual adenine bases over a ±250 bp window, comparing 6mA-modified adenines against all genomic adenines, computed in 10 bp bins with a smoothening window of 3. Additionally, 6mA density was quantified across nucleosome and linker DNA regions, with nucleosome positions divided into 10 bins and linker regions into 4 bins (linker length = 200 bp). This analysis was performed within a ±500 bp window around nucleosome dyad positions divided into 100 bins, with flank normalisation applied over 300 bp flanking regions and a smoothing window of 7. Up to 50,000 6mA sites per timepoint were included in each analysis.

### TSS analysis

6mA site density was profiled around annotated TSSs derived from 5’UTR features in the PlasmoDB-68_Pfalciparum3D7.gff annotation. Using a custom Python script (plot_tss_profiles.py), 6mA density was computed over a ±1,000 bp window centred on each TSS, divided into 100 bins with a smoothing window of 5, and reported as 6mA density per kb. This analysis was performed separately for all annotated genes and for the subset of genes carrying at least one 6mA modification, across all four timepoints. For the 20h timepoint, deepTools was additionally used to generate TSS correlation plots over a ±500 bp window.

### RNA-seq and data analysis

To investigate the relationship between 6mA modification and gene expression, steady-state transcript levels were obtained from a publicly available RNA-seq dataset [49] and processed to reads per kilobase pair per million (RPKM). Genes were first divided into two broad groups: those carrying at least one 6mA modification and unmodified genes. Modified genes were further stratified by the genic location of their 6mA sites into six categories: 5’UTR only, CDS only, 3’UTR only, 5’UTR and CDS, 3’UTR and CDS, and 5’UTR, CDS and 3’UTR. Expression levels across all groups were compared and visualised using ggplot2 with statistical significance between groups assessed using a Wilcoxon test [81].

## Acknowledgements

This work was supported by funding from the Department of Electronics, IT, BT & ST, Government of Karnataka, to IBAB, Bengaluru, the Ramalingaswami Re-entry Fellowship from the Department of Biotechnology, Government of India, to SSV (BT/RLF/Re-entry36/2017) and a SERB-CRG grant from the Department of Science and Technology, Government of India, to SSV (CRG/2020/004801). SD is supported by a National Eligibility Test-Senior Research Fellowship (NTA Ref No.: 191620123696) awarded by the University Grants Commission, India. SMRT sequencing on the Revio system performed at the Sequencing Technology Center at the University of Louisville and was supported by NIH S10 OD034432 awarded to MLS. Work in the Greer lab is supported by NIH R01AG084739 and NSF award number 2326672.

## Authors’ Contributions

D.S. contributed to data acquisition, formal analysis, investigation, methodology, validation, visualization, data curation, and writing of the original draft and subsequent revisions. W.L. contributed to data acquisition (SMRT sequencing), investigation, methodology, and manuscript review and editing. G.S. contributed to data acquisition (Fig. 2A), formal analysis, investigation, methodology, and manuscript review and editing. M.G. contributed to formal analysis, investigation, and manuscript review and editing. C.S.M. contributed to formal analysis, methodology, and manuscript review and editing. E.L.G. contributed to formal analysis, methodology, funding acquisition, project administration, and manuscript review and editing. M.L.S. contributed to formal analysis, methodology, funding acquisition, project administration, and manuscript review and editing. S.S.V. contributed to conceptualization, formal analysis, funding acquisition, investigation, methodology, project administration, supervision, validation, visualization, and writing of the original draft and subsequent revisions. All authors read and approved the final manuscript.

## Data availability

The raw data generated by SMRT sequencing and RNA-seq fastq files have been deposited in the NCBI SRA Repository under the Accession No. PRJNA1475858 (http://www.ncbi.nlm.nih.gov/bioproject/1475858).

