## Supplementary material for "DNA 6mA marks transcriptionally active chromatin in malaria parasites": SD_20260525_Supplementary Data.docx

**Supplementary Information for Seshan et al., 2026**

This document contains:

**Supplementary Tables S1 and S5**

**Figures S1-S9**

The following supplementary tables are provided as excel sheets:

**Supplementary Tables S2: 6mA sites considered for downstream analysis across all timepoints**

**Supplementary Tables S3: List of 942 genes that are modified with 6mA across all timepoints**

**Supplementary Tables S4: RNA-seq read counts (raw and normalised) for 10 h replicates**

**Supplementary Tables S6: SMRT-seq and RNA-seq statistics**

**Supplementary Table S1: List of known 6mA DNA Methyltransferases used as BLASTp queries against the *P. falciparum* proteome**

| **Enzyme** | **Organism** | **Name of Protein** | **Source (Reference)** | **UniProt ID** |
| --- | --- | --- | --- | --- |
| **6mA Methyl-transferase** | *Homo sapiens* | N6AMT1 | (Xiao et al., 2018) | Q9Y5N5 |
|  | *Tetrahymena thermophila* | TAMT - 1 | (Luo et al., 2018) | Q23RE0 |
|  | *Oxytricha trifallax* | MT-A70 | (Beh et al., 2019) | J9IF92 |
|  | *Caenorhabditis elegans* | DAMT-1 | (Greer et al., 2015) | Q09956 |
|  | *Mus musculus* | METTL4 | (Zhang et al., 2020) | Q3U034 |

**Supplementary Table S5: List of oligonucleotides used in this study**

| **Name** | **Sequence (5' to 3')** | **Description** |
| --- | --- | --- |
| SD01 | GATCGGATCCATGAAAAATTCTGAACAAGTCAG | Forward primer for amplifying the PfN6AMT gene from *P. falciparum* genomic DNA |
| SD02 | TGCCTCGAGTTATTTTTTCTTCATTAATTTGTAAATAAATATTGTTTCG | Reverse primer for amplifying the PfN6AMT gene from *P. falciparum* genomic DNA |
| SD03 | ATATCGGAATTAATTCGGATCC C ATGGCAGGGGAGAACTTCG | Forward primer for amplifying the hN6AMT1 coding sequence from human cDNA |
| SD04 | GTGGTGGTGGTGGTGCTCGAGAGACTTGGTGAACTTGAGGAC | Reverse primer for amplifying the hN6AMT1 coding sequence from human cDNA |
| SD05 | TTTGATATTGTTTTATTTGCTGCAGCAGCTGTTATTACAGGACCAGATG | Forward primer for creating catalytically dead version of the PfN6AMT gene |
| SD06 | CATCTGGTCCTGTAATAACAGCTGCTGCAGCAAATAAAACAATATCAAA | Reverse primer for creating catalytically dead version of the PfN6AMT gene |
| SD07 | AAGTATATAATATTGGTGAAGGTGATGCAACATA**G**TTTTAGAGCTAGAA | DNA oligonucleotide used in methyltransferase activity assay (forward); annealed with SD08 to create a dsDNA fragment that was used in the assays |
| SD08 | TTCTAGCTCTAAAA**C**TATGTTGCATCACCTTCACCAATATTATATACTT | DNA oligonucleotide used in methyltransferase activity assay (reverse) |

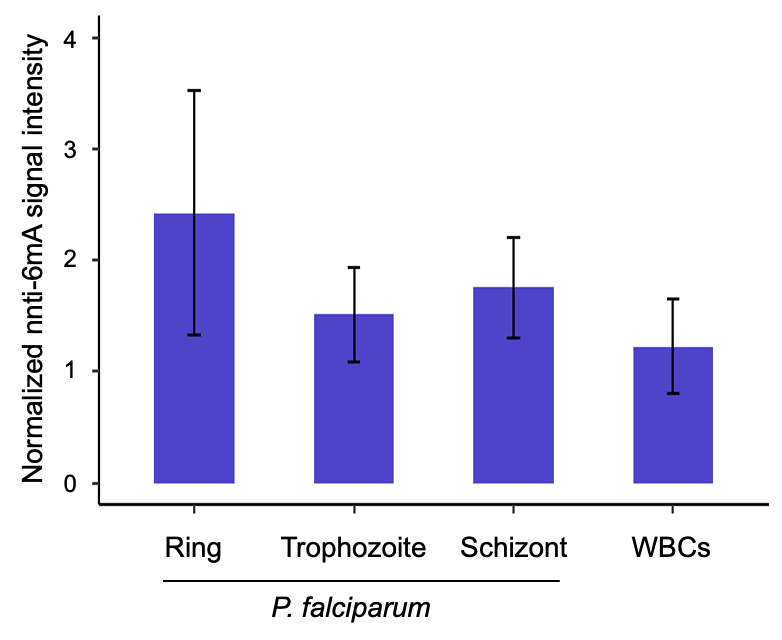

**Supplementary Figure S1:** **Quantification of 6mA levels in *P. falciparum* genomic DNA, as detected by South-western blotting with anti-6mA antibodies.** Y-axis represents the signal intensity of anti-6mA antibody reactivity with genomic DNA normalized to the amount of genomic DNA loaded (*i.e.,* methylene blue signal intensity). Data represent the mean ± standard deviation of a minimum of three biological replicates.

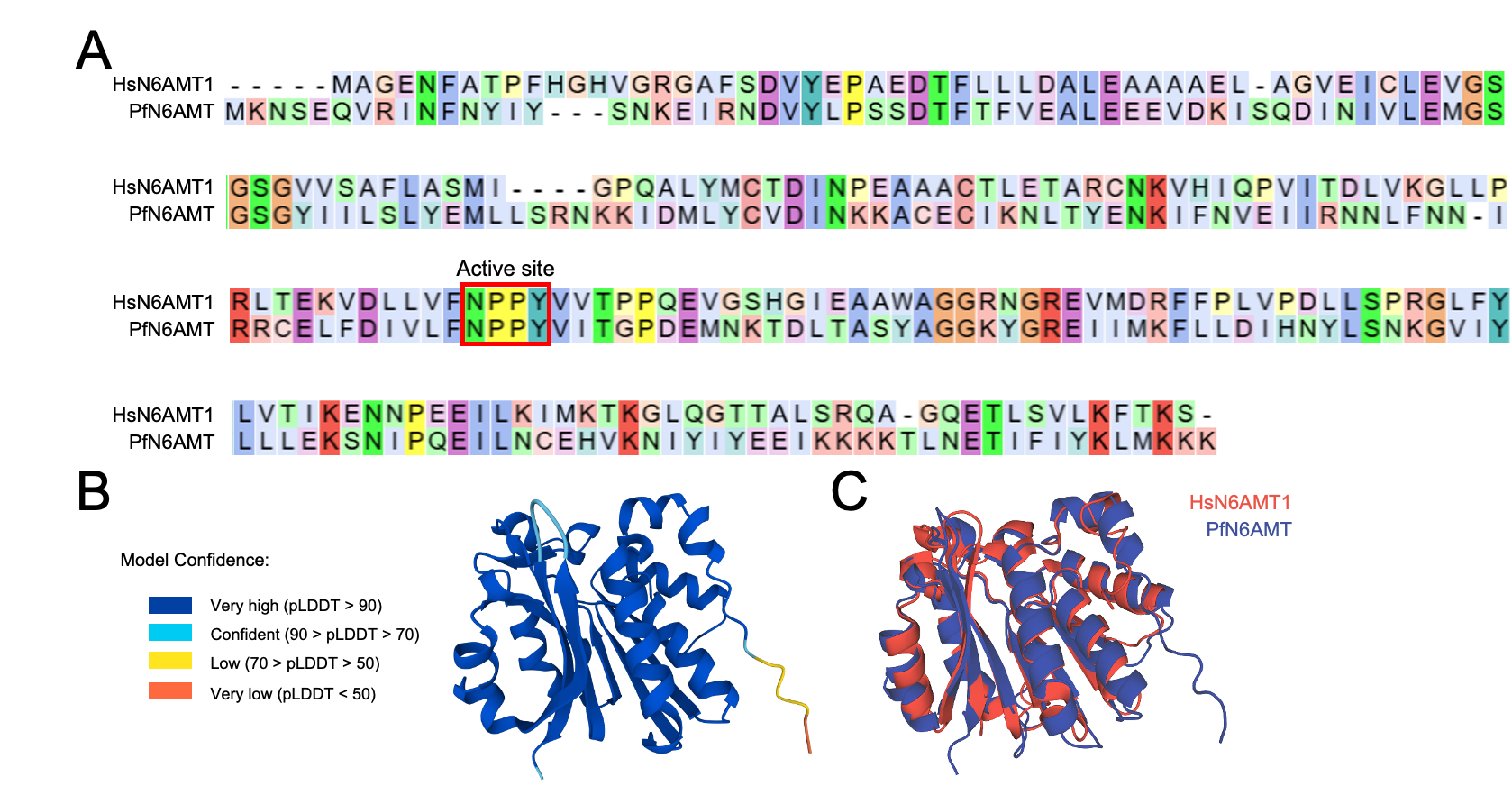

**Supplementary Figure S2: *In silico* analysis of the putative *P. falciparum* 6mA methyltransferase PfN6AMT**. ***(A)*** Pairwise sequence alignment of human N6AMT1 (Uniprot ID: Q9Y5N5) and PfN6AMT (*PF3D7_1303100*) that was identified using BLASTp. The active site residues for the human protein are highlighted (NPPY at positions 122 to 125). ***(B)*** The AlphaFold-predicted structure of PfN6AMT (https://alphafold.ebi.ac.uk/entry/Q8IET4). ***(C)*** Structural overlay of the AlphaFold-predicted structure of PfN6AMT onto the crystal structure of human N6AMT1 (PDB ID: 6KMR; Li et al., 2019), visualized using PyMol (DeLano *et al.,* 2002). RMSD of all residues is 1.6 Å.

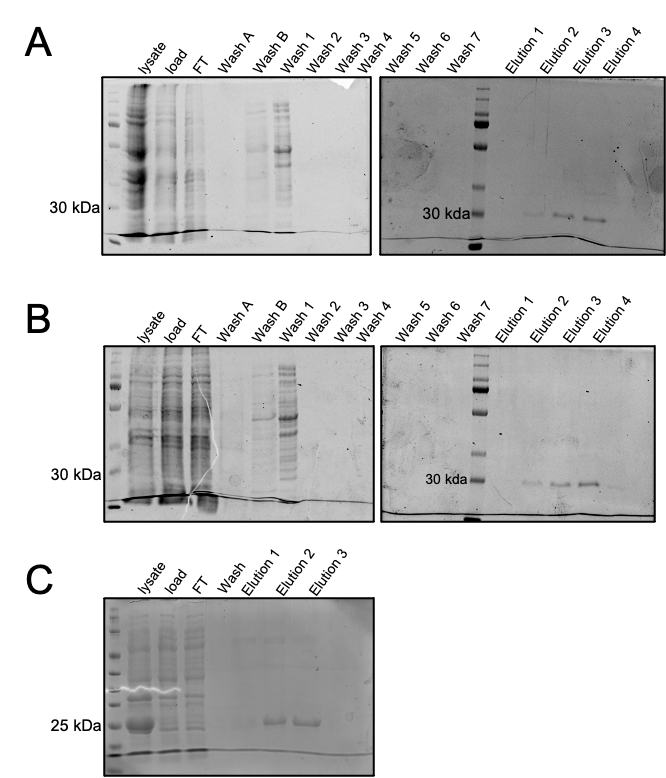

**Supplementary Figure S3: Purification of 6XHis-tagged recombinant proteins using Ni^2+^-NTA affinity chromatography.** Fractions collected during the purification of ***(A)*** wild-type PfN6AMT, ***(B)*** catalytically dead PfN6AMT, and ***(C)*** hN6AMT1 were resolved using SDS-PAGE and visualised using Coomassie Brilliant Blue staining. FT = Flowthrough.

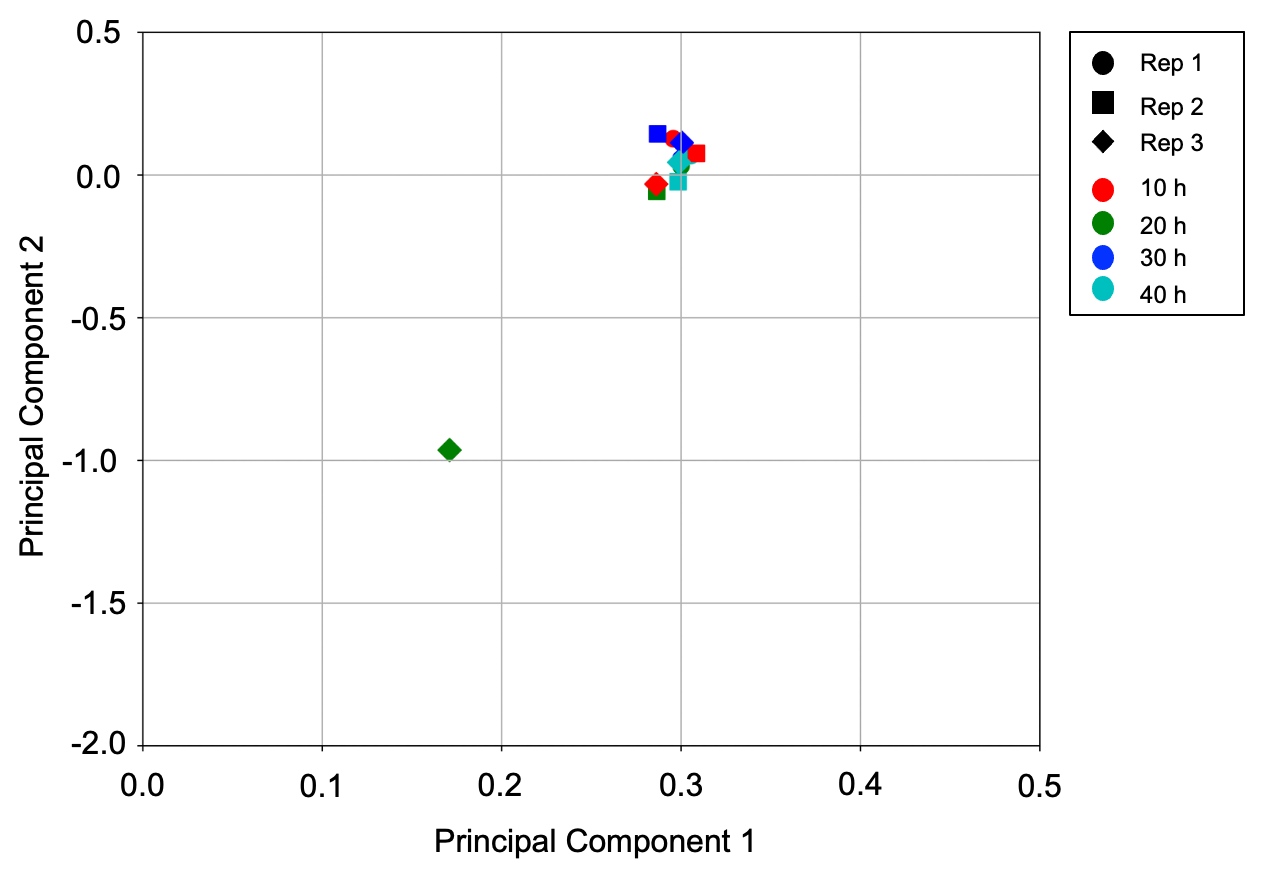

**Supplementary Figure S4:** **Principal Component Analysis of SMRT-seq-derived 6mA genome-wide profiles for the 12 *P. falciparum* samples sequenced in this study.** Principal Components 1 and 2 capture 74.89 % and 6.52 % of the variance, respectively.

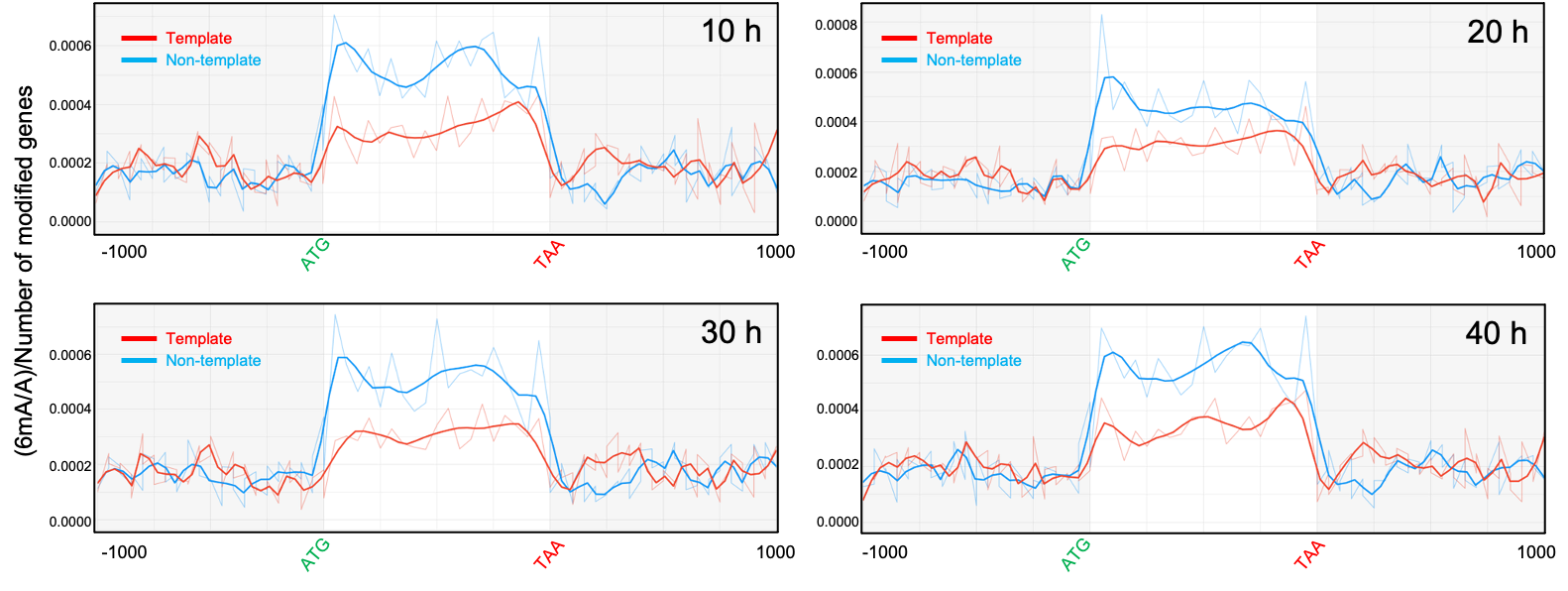

**Supplementary Figure S5: 6mA modification levels for template and non-template strands of modified genes at four timepoints of *P. falciparum* IE development.** The gene body (ATG to TAA, including introns) and 1000 bp upstream/downstream flanking regions of modified genes were divided into 20 equal-sized bins and the 6mA/total A ratio calculated for each bin for the template (red) and non-template (blue) strands. The Y-axis represents the average value across all modified genes at that timepoint.

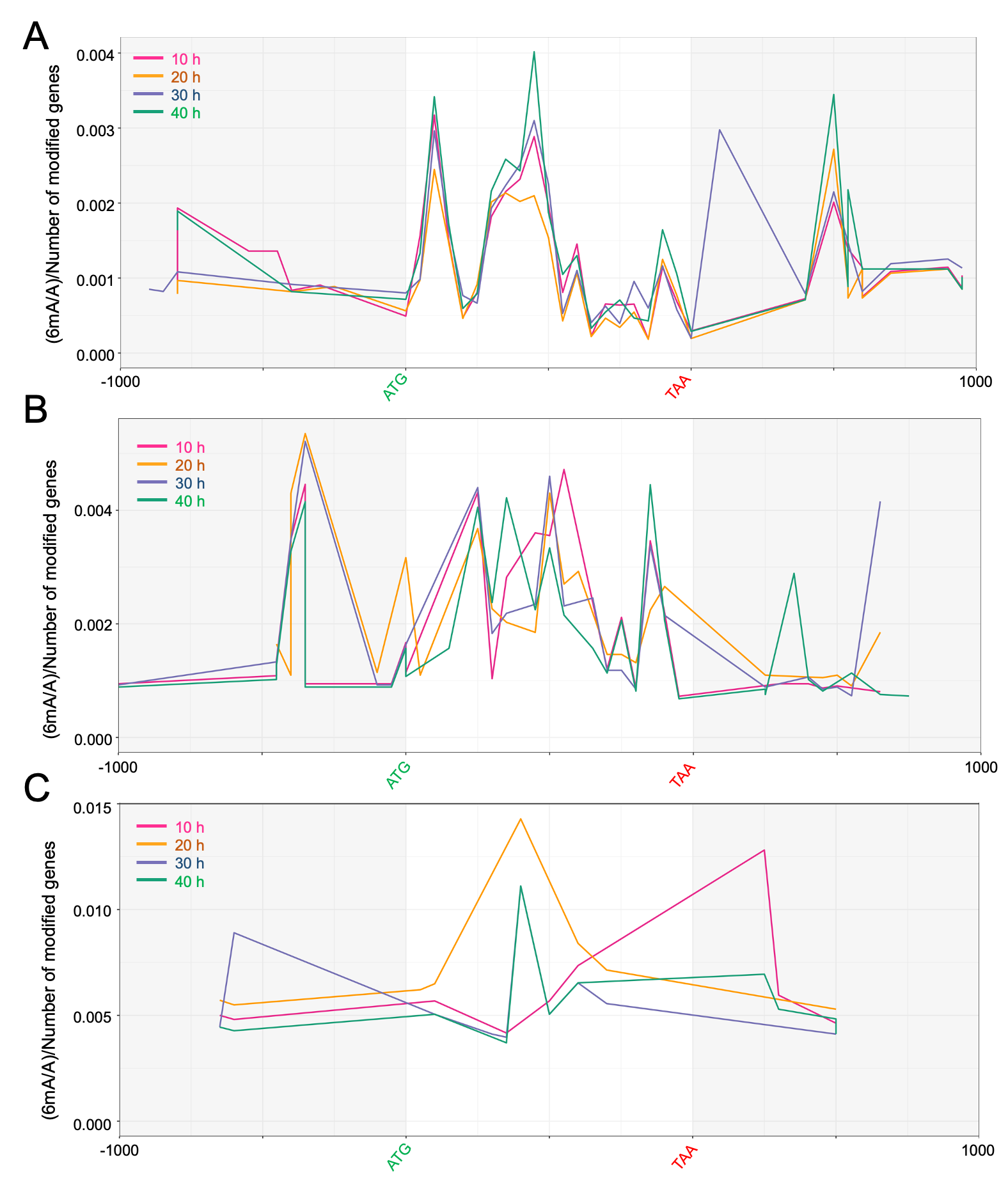

**Supplementary Figure S6: Comparing the 6mA profiles of *var, rifin* and *stevor* gens across different timepoints.** 6mA levels across the start codon (ATG), gene body, stop codon (TAA) and 1000 bp upstream and downstream flanks of ***(A)*** *var,* ***(B)*** *rifin* and ***(C)*** *stevor* multigene families. Each region was divided into 20 equal-sized bins for standardized representation.

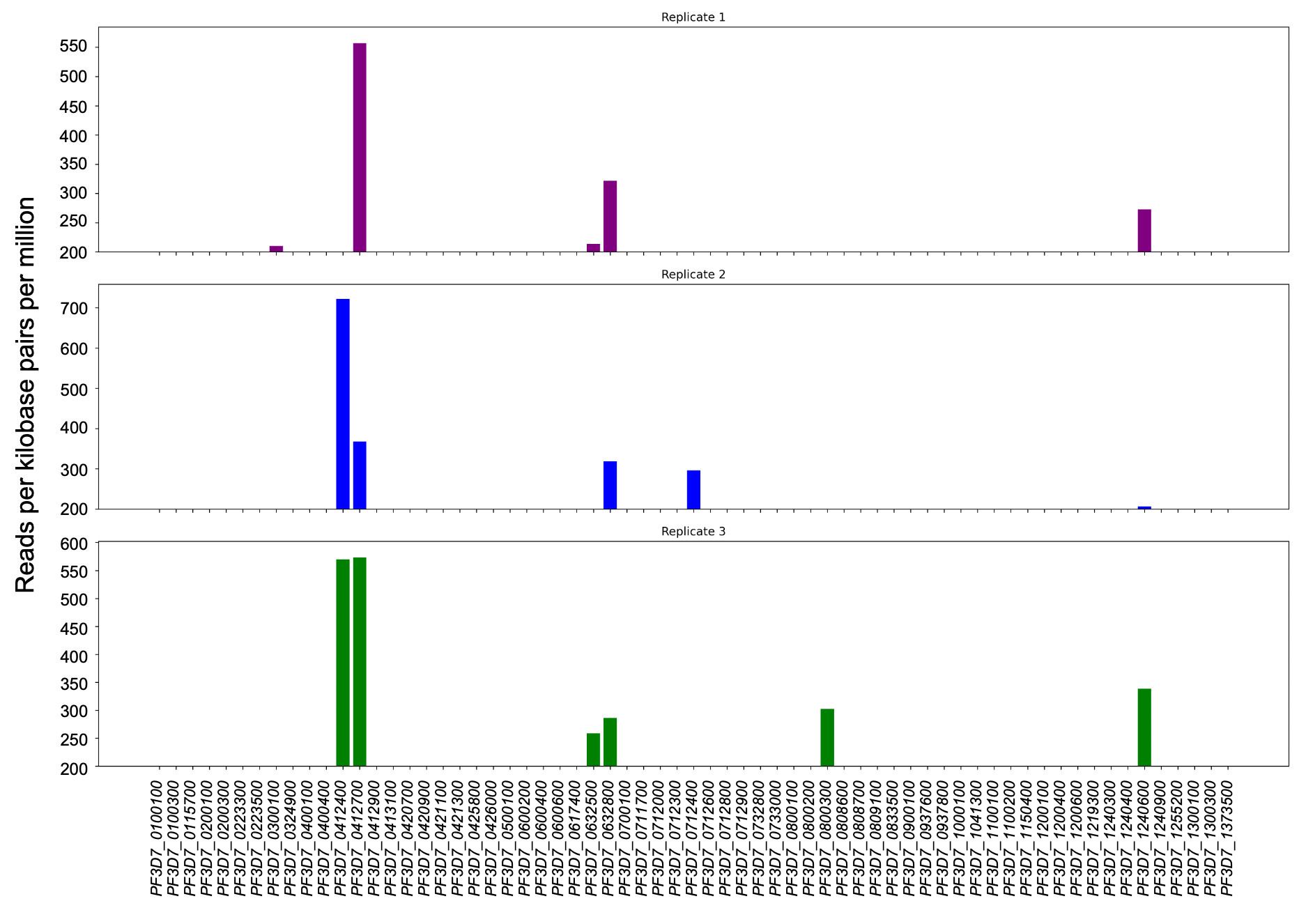

**Supplementary Figure S7: Expression analysis of *var* genes in the 10 h SMRT-seq samples.** Strand-specific RNA-seq was performed on the 10 h *P. falciparum* 3D7 samples that were harvested for SMRT-seq. Normalised RNA-seq read counts for all *var* genes is shown for the three replicates.

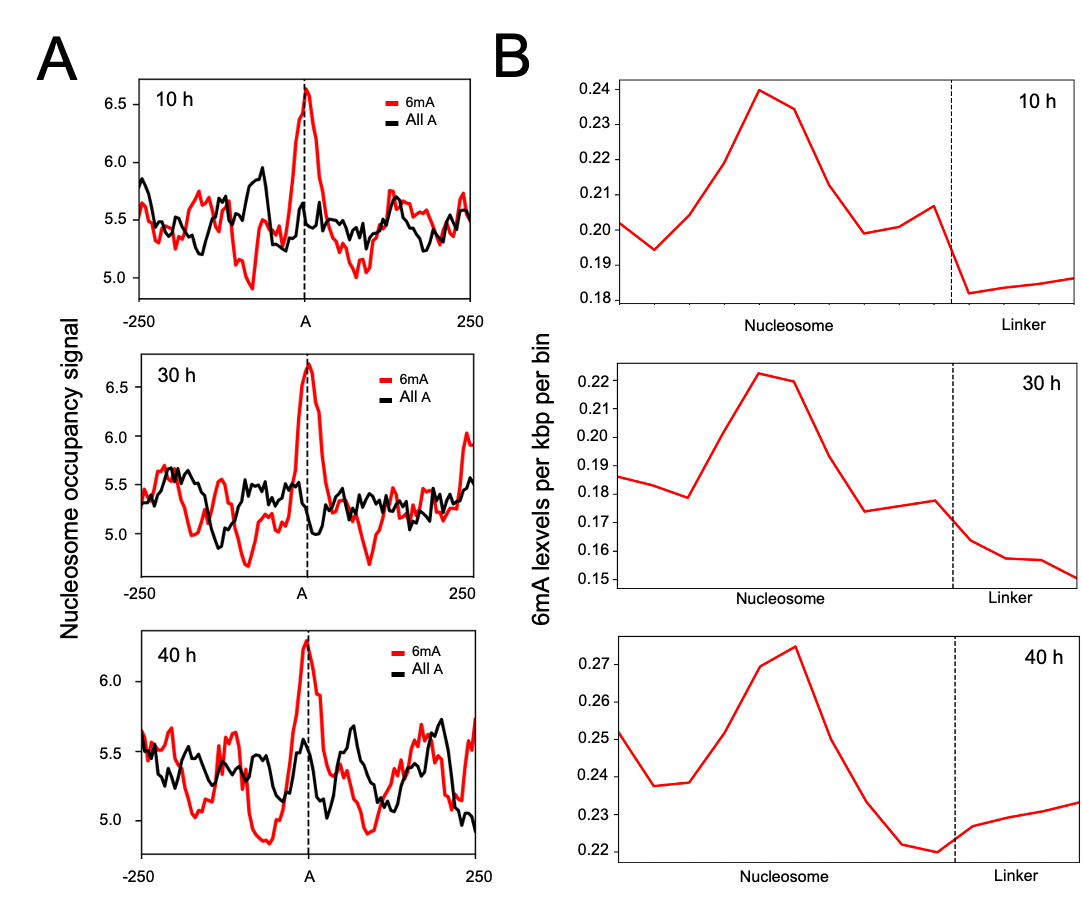

**Supplementary Figure S8: Correlation analysis between 6mA and nucleosome occupancy (Kensche et al., 2016) at 10, 30 and 40 h timepoints.** ***(A)*** The X-axis represents an adenine base anchored at “0” and its +/- 250 bp flanking regions. The Y-axis region represents the nucleosome occupancy signal. The red line is centred at 6mA loci while the black line is centred at an equal number of adenosine sites chosen at random. ***(B)*** The X-axis represents the position of the nucleosome (divided into 10 bins) and linker (divided into 4 bins). The Y-axis represents 6mA levels per kbp per bin.

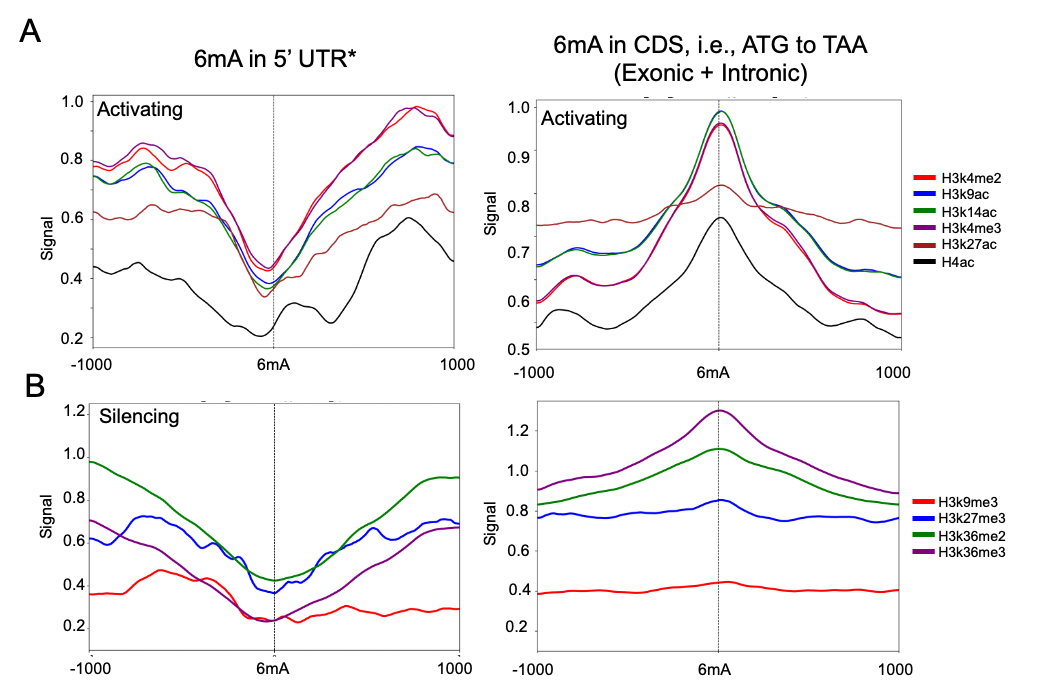

**Supplementary Figure S9: Correlation analysis between 6mA and histone PTMs at 40 h. *(A & B)*** DeepTools plotprofile was used to compare the read density distribution of the indicated activating ***(A)*** and silencing **(B)** histone PTMs (Bártfai et al., 2010; Karmodiya et al., 2015) for all 6mA positions located within 5’UTRs (*left*) or gene bodies (*right*) of modified genes at 40 h.

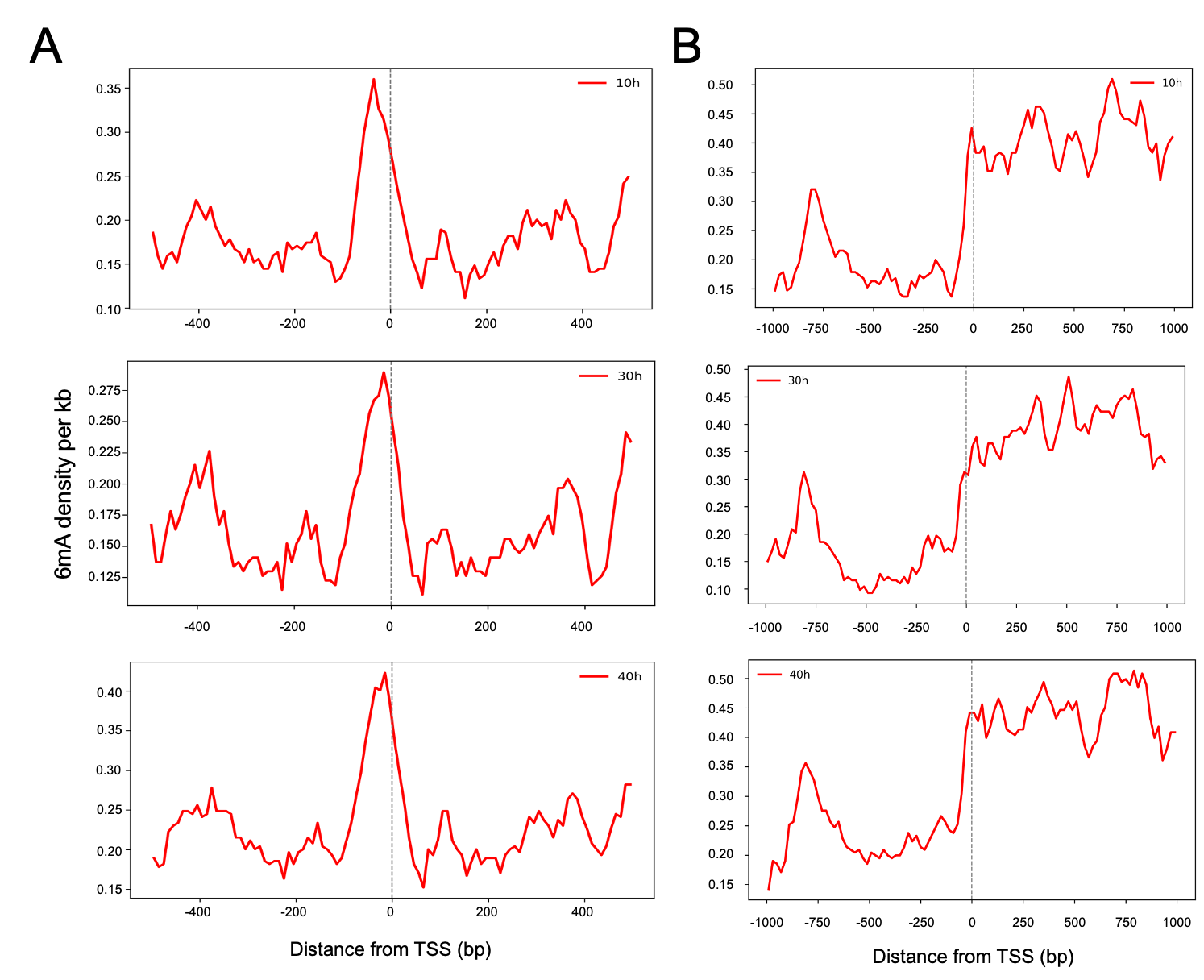

**Supplementary Figure S10: Density distribution of 6mA around TSS at 10, 30 and 40 h.** DeepTools was used to analyse the density of 6mA +/-500 bp from TSS (https://plasmodb.org) for 10 h, 30 h and 40 h data. ***(A)*** All TSS, and ***(B)*** TSS of modified genes alone.

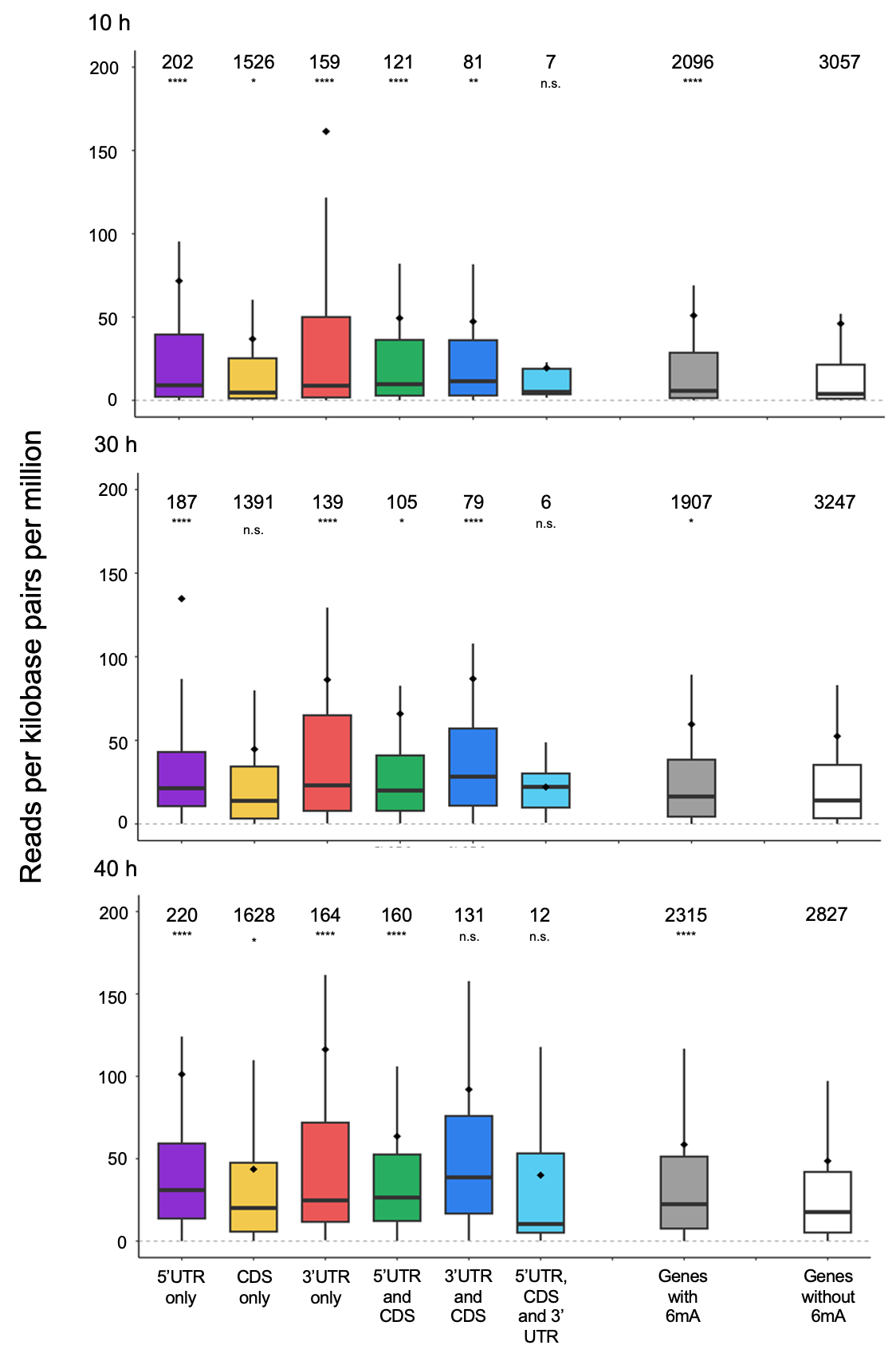

**Supplementary Figure S11: Correlation analysis between 6mA in genic features and steady state transcript levels.** Box plots were used to represent the distribution of normalised RNA-seq read counts (Reads per kilobase per million mapped reads (RPKM)) for 6mA-modified genes at the 10 h, 30 h and 40 h timepoints. RNA-seq data were obtained from (Kensche et al., 2016). The box shows the interquartile range, with the line inside representing the median; the mean is indicated using a  ♦ . The numbers of genes in each group are shown. p-values were calculated using a Wilcoxon test. UTR = Untranslated region; CDS = Coding sequence.
